# Is earlier reproduction associated with higher or lower survival? Antagonistic results between individual and population scales in the blue tit

**DOI:** 10.1101/2021.01.11.426202

**Authors:** Olivier Bastianelli, Anne Charmantier, Clotilde Biard, Suzanne Bonamour, Céline Teplitsky, Alexandre Robert

## Abstract

Although it has been shown that phenology can respond to temporal environmental variation in free ranging populations of several species, little is known about the mechanisms of these responses and their effects on demography, and in particular on survival. Exploring phenological responses and their associated consequences on survival can be achieved at two distinct scales: the population scale, which focusses on a set of common responses to environmental conditions, and the individual scale, focusing on the relative position of each individual in the distribution of survival and phenology under particular conditions. In this study, we apply capture-mark-recapture multistate modelling on a 38-year monitoring dataset of blue tits (*Cyanistes caeruleus*) to investigate the effects of breeding phenology and some demographic covariates (breeding density, average and individual breeding success) on adult survival, at both population and individual scales. Our analysis revealed that (i) at the population scale, early breeding years are followed by lower average adult survival. (ii) At the individual level, earlier breeders within the population have higher subsequent survival than later breeders, although this relationship is reversed in years with very harsh conditions, e.g. warm spring and high breeding density. (iii) High individual relative breeding success is also associated with higher subsequent survival and explains more survival variation than relative phenology. Overall, our study indicates that, although earlier breeding is associated with a survival cost at the population level, substantial intrapopulation hererogeneity shapes a positive association between earlier breeding, breeding success and survival at the individual level.

## INTRODUCTION

The phenological response of organisms to variation in their local environment is a major issue in evolutionary ecology and conservation research, both because the diversity of this response reflects differences in evolutionary potential and strategies between and within species (Thackeray et al., 2016), and because advances in reproductive phenology is seen as one of the universal ecological responses to global warming (Durant et al., 2007; Parmesan, 2007; Radchuk et al., 2019). Although it has been shown that phenology can respond to meteorological variations (noise and cycles) as well as climatic trends in many plant (Bjorkman et al., 2015; Diekmann, 1996) and animal (Duursma et al., 2018; Samplonius et al., 2018; Sheriff et al., 2015; Haest et al., 2018) species, little is known about the mechanisms of these responses and their effects on demography and population dynamics (Forrest and Miller-Rushing, 2010, but see Shipley et al., 2020; Simmonds et al., 2020).

In particular, it is likely that phenological variations, which may involve variations in strategy within populations, are constrained by trade-offs (Diggle, 1999; Forrest and Miller-Rushing, 2010). For example, in birds, while early reproduction might be beneficial in terms of reproductive success (Harriman et al., 2017; in the focal population: Marrot et al., 2018), it is also potentially associated with subsequent survival cost for the breeders (e.g. Brinkhof et al., 2002; Nilsson, 1994) or the early hatched nestlings (Shipley et al., 2020). The presence of these trade-offs could partly explain phenological mismatches (*i.e*. a difference between a species phenology and its main resource phenology) frequently observed in bird populations (Visser et al., 2012). Although a review of the literature provides multiple estimations of reproductive selection acting on avian breeding phenology (Verhulst and Nilsson, 2008; Charmantier and Gienapp, 2014), similar links between survival and timing of breeding are too scarce to conclude on the presence of such trade-offs. One reason why survival has not been explored frequently in this context might be because it is often assumed that reproductive success will be affected by breeding phenology while survival will be influenced by migration phenology (e.g. Visser and Gienapp, 2019).

Additionally, the survival consequences of phenological variations may be influenced by other mechanisms such as intraspecific competition. Several ecological field studies (in both long-lived and short-lived species) have reported lower population level survival rates at high densities (Abadi et al., 2012; Fay et al., 2015; Brouwer et al., 2006; Bonenfant et al., 2009; Le Cœur et al., 2016) and competition has been shown to influence population dynamics in many studies (see e.g. Grøtan et al., 2009 and Gamelon et al., 2016 in the Great tit *Parus major*). Importantly, however, in the presence of both year-to-year environmental changes and intraspecific competition, population density can also be a factor maintaining demographic stability in a changing environment. Reed and colleagues (Reed et al., 2013) showed that in a Great tit population, the reduced reproductive success observed in years with greater mismatch between the tits and their prey was compensated by a higher overwinter survival due to relaxed competition.

Overall, exploring breeding phenological responses and their associated demographic benefits and costs can be achieved at two distinct scales: a/ the population scale, by studying for example the overall response of a population (e.g. mean or median phenology, reproductive success or survival) to interannual environmental variation, or b/ the individual level, by focusing on the relative position of each individual in the distribution of reproductive success, survival and phenology under particular conditions (e.g. for the same year, see e.g. Harriman et al., 2017 for an experimental approach focusing on breeding success). The first approach examines a set of common responses (behavioural, demographic) to environmental conditions, while the second approach focuses on individual heterogeneity within a population, in terms of fitness components. While some external constraints induced by the limitation of resources for survival and reproduction are shared by all individuals of the population, differences can arise in resource acquisition due to e.g. genotype or developmental conditions (Lindström, 1999; van Noordwijk and de Jong, 1986; Wilson and Nussey, 2010).

In the context of exploring demographic consequences of phenological changes, investigating variation within and between individuals is of primary importance as it occurs in virtually all traits, including demographic components such as reproduction and survival (Clutton-Brock, 1988). Between-individual variation may be linked with phenotypic attributes that change along an individual’s life, such as age or breeding status, or with fixed attributes (such as sex or genotype, see Gimenez et al., 2018 and references therein). A significant part of the research on individual heterogeneity has focused on the covariation between demographic components such as reproduction and survival. In particular, a negative relationship between survival and reproduction is expected in the context of a trade-off (energetic or genetic) between these two components (Harshman and Zera, 2007; Roff, 1993; Stearns, 1992). On the contrary, a positive relationship is expected within the framework of dynamic (Tuljapurkar et al., 2009) and/or fixed (Cam et al., 2013) variations between individuals. Fixed individual heterogeneity is linked, for example, to intrinsic or resource acquisition differences early in life (Hamel et al., 2009; Lescroël et al., 2009), sometimes referred to as individual quality (Cam et al., 2002; Wilson and Nussey, 2010). Reproductive costs and individual quality are not mutually exclusive and have been suggested to interact with each other as well as with environmental (Robert et al., 2012) and individual (Robert et al., 2015) conditions.

Although phenological variation within populations has been well described in many organisms (Elzinga et al., 2007; Forrest and Miller-Rushing, 2010; Laaksonen et al., 2006), the relationship between such variation and demographic heterogeneity remains poorly understood both empirically and theoretically. Even in the well-studied context of temperate forest passerines where early seasonal breeding has been repeatedly associated with higher breeding success, the theoretical frameworks outlined above on trade-offs *versus* resource acquisition and individual heterogeneity can lead to two opposing predictions. First, one could predict that, in a given population a given year, the earliest breeders will show the lowest survival due to a reproduction-survival trade-off (Brinkhof et al., 2002; Wiggins et al., 1998; Nilsson, 1994; Nilsson and Svensson, 1996). Alternately, one could expect that the high “quality”, earliest breeders (e.g. with highest resource acquisition, van Noordwijk and de Jong, 1986), will exhibit the highest survival. Additionally, the question remains whether the observed phenology-survival pattern is consistent across years, in particular whether it is similar in years with early or late average breeding dates. Answering these questions requires studying the links between reproductive phenology and survival, first at the population level, second at the individual heterogeneity level, and third the links between these two levels.

In this study, using a long-term monitoring dataset, we investigated the effect of breeding phenology on adult survival in a population of a temperate passerine bird, the Blue tit (*Cyanistes caeruleus*). The data comprised 1562 adult individuals captured and ringed in nest boxes during the reproduction period across 38 years (1979-2016). In this population, breeding phenology is influenced by meteorological conditions. Median laying date is earlier in warmer years (Bonamour et al., 2019) and has an impact on breeding success: earlier breeding is correlated with higher number of fledglings at the individual and population scales (Marrot et al., 2018). Long-term effects of climate on the phenology of the studied population also occurred over the period of the study (Marrot et al., 2018), during which spring temperatures have significantly increased (1979-2016: +0.49°C/decade; *p-value* < 10^-3^) and median laying dates have significantly advanced (1979-2016: −2.97 days/decade; *p-value* < 10^-3^).

Using flexible capture-mark-recapture multistate modelling (Lebreton et al., 2003), we addressed four different questions: First, is there a link between population-level phenology and average annual adult survival in the focal Blue tit population? Since previous experimental studies in birds (Brinkhof et al., 2002; Wiggins et al., 1998) including blue tits (Nilsson, 1994; Nilsson and Svensson, 1996) have revealed a cost of early breeding on adult survival, we predict a low average adult survival following early population breeding phenology. Second, within a given year, what is the link between individual timing of breeding (quantified as laying date quartiles of the population for this particular year, hereafter *relative laying date*) and adult survival? We expect to find either a positive association between early breeding and survival if “good quality” individuals are those that invest in early breeding (Robb et al., 2008) or, on the contrary, a negative association if the cost of breeding earlier outweighs differences in individual heterogeneity (Tarwater and Arcese, 2017). Third, does the link between the relative laying date and adult survival vary depending on the conditions faced in a given year (e.g., population density)? Here we expect to find a weaker link between relative laying date and adult survival in years with low constraints on breeding (e.g. low population density) and therefore probable relaxed cost of breeding. Fourth and finally, since breeding early and having a high breeding success are correlated (*i.e*. early broods are larger: review in Klomp, 1970, in the focal population Delahaie et al., 2017) but may convey distinct costs or reflect distinct facets of individual heterogeneity, our last interrogation is about the link between relative laying date, survival and individual breeding success. Specifically, are large brood sizes relative to the population associated with high or low survival, and what is the magnitude of this effect relative to the effect of laying date?

## Methods

### 1 Population Monitoring

Our data was collected over 38 years (1979-2016) in a natural population of blue tits *Cyanistes caeruleus* in an evergreen oak (*Quercus ilex*) forest on the Mediterranean island of Corsica (E-Pirio site: Lat 42.38; Long: 8.75; see Blondel et al., 2006 and Charmantier et al., 2016 for details). Blue tits are cavity-nesting non-migratory passerine birds that readily breed in nest boxes, which they often prefer to natural cavities (Newton, 1994). The high density of nest boxes compared to the rarity of natural cavities in the E-Pirio site (A.C., unpublished data) allowed us to monitor almost all breeding individuals in the E-Pirio study area. During the whole study period, nest boxes were checked every year at least weekly during the breeding season (March to June), allowing to accurately assess the date of the first laid egg for all clutches (hereafter, laying date), as well as the clutch size and the number of fledglings. Breeders were captured in the boxes when nestlings were 10-15 days old, *i.e*. late enough to avoid the risk of nest abandonment, but early enough to avoid early departure by the nestlings. Parents were identified with a unique numbered metal ring provided by the Centre de Recherches sur la Biologie des Populations d’Oiseaux (CRBPO, which also provided the permits under which capture and handling of birds were conducted). Sex and age at first capture were determined by plumage patterns for every breeder. As these patterns cannot be used to age individuals older than two years, only two adult age classes were used in survival modelling: 1 year and 2 or more years. Individuals whose age at first capture had not be assessed were removed (3.4% (n=55) of the individuals). In total, 1562 reproducing individuals were considered in this study. All data analysed concerned first broods only (second broods represent less than 2.5% of breeding events in this population).

### 2 Population-level annual covariates

For each year of the study, we computed the median of individual laying dates in the population (median laying date, medLD), the mean number of fledglings per attempted first brood in the population (reproductive success, repS) and the arcsine normalized breeding density of the population (population density, popD).

Population breeding density was obtained by measuring nest box occupation rates by blue tits in a defined area of the study site, where nest boxes number varied as little as possible over the 38 years of study (n=58 to 62 nest boxes). Only blue tit occupation was considered as the diameter of the nest boxes entry hole (26 mm) was too small to allow regular occurrence of Great tit (*Parus major*) nesting.

The variable used to study how meteorological conditions affect individual survival and reproduction was the average daily mean temperature during the period from March 31^st^ to May 7^th^ (hereafter, Temp). This 38 day window has been identified in a previous study using a sliding window analysis, as the time window most correlated with the average annual laying date in the focal population (see Table 1 in Bonamour et al., 2019). The temperature data were obtained by regressing daily temperature measurements over 4 years in the EPirio station with temperature data from the meteorological station of Calvi (Lat: 42.52; Long: 8.79, 15.9 km away from the population) and inferring the temperature in E-Pirio over the remaining years.

**Table 1:** Summary of the four main types of questions addressed and capture-mark-recapture models used in this study.

|  | Population-level analysis | Individual-level analysis | Additional annual population-level covariates tested on survival | Type of models used |
| --- | --- | --- | --- | --- |
| a) Link between population-level phenology and average annual adult survival | Phenology ↔ Survival |  | Demographic covariates<br>(median laying date, population density, mean reproductive success) | Monostate models |
| b) Link between individual timing of breeding (relative laying date) and adult survival |  | Relative Phenology ↔ Survival |  | Multistate models<br>(4 relative laying date states) |
| c) Effects of population-level annual covariates on the relative laying date - adult survival link |  | Relative Phenology ↔ Survival | Demographic covariates<br>and spring temperature | Multistate models<br>(4 relative laying date states) |
| d) Links between individual breeding success, relative laying date and adult survival |  | Relative Laying date and<br>Relative Reproductive Success<br>↔ Survival |  | Multistate models<br>(6 relative laying date and<br>reproductive success states) |

### 3 Capture-Mark-Recapture modelling

Individual capture-recapture histories for the 1562 breeding individuals were analysed to provide robust estimates of survival and recapture probabilities (respectively *φ* and *p*, Lebreton et al., 1992), using a logit-link function. Monostate Capture-Mark-Recapture modelling was used for the assessment of the link between annual population-level phenology and annual survival, while multistate capture-mark-recapture models (Lebreton et al., 2003) were necessary for assessing the link between intra-population variation of phenology (i.e., relative laying date), breeding success and adult survival. Multistate models make it possible to model the survival of individuals that may be in different states (which correspond to different categories of laying date or reproductive success in the case of our study). These models include four types of parameters: the probabilities of being in the various initial states (IS), the probabilities of transition among states (T), survival probabilities (*φ*) and recapture probabilities (*p*). All analyses were conducted using the program E-SURGE (Choquet et al., 2009b). Goodness-of-fit of models to the data was ensured for each dataset using the program U-CARE (Choquet et al., 2009a), based on the Cormack Jolly Seber model for monostate models and on the Jolly Move model for multistate models.

In several years, experiments were conducted in the population including brood size manipulations that can highly alter adult survival (Dijkstra et al., 1990; Nur, 1984). Capture-recapture histories of the corresponding individuals were censored right after the first time they experienced such a fitness-changing experiment. However, nestlings born during these experiments and later recaptured as adults were not removed from the analysis. This necessary censoring resulted in a significant decrease in observation numbers (−23.6% from 2955 to 2258 capture and recapture events in the monostate dataset). Breeding events in experimental conditions were also removed from reproductive success estimations.

### 4 Data analysis and model selection

The Akaike’s Information Criterion corrected for small sample size (AICc) was used for model selection (Burnham and Anderson, 1998). A low AICc was considered revealing a good compromise between the fit to the data (likelihood of the model) and the number of parameters used by the model. The threshold for significant difference between two models was set at two AICc points. In case of lower difference, the model with the lowest number of parameters was selected. The four main questions addressed are summarised in Table 1 and details are provided in the sections below.

#### a Link between population-level phenology and average annual adult survival

Prior to assessing the effects of population-level covariates on survival, a preliminary model selection assessed the effects of year, sex and age on recapture (*p*) and survival (*φ*) probabilities with the monostate dataset of capture histories (Table 2A). The starting model (Model 22) included potential effects of sex, age and yearly variations (hereafter, *year*) on survival and potential effects of sex and year on recapture probabilities. Only simple (first order) interactions between variables influencing survival and recapture probabilities were considered, to enable robust biological interpretation of the results and avoid overcomplexity. All subsequent models were nested in this one. The model selection started with simplifying constraints on *p*. As a second step, we retained the best model structure for recapture and considered simpler survival models. We then used the best model to test the effects of our three demographic temporal covariates (medLD, popD, repS) and their interactions (Table 2B) on survival.

#### b Link between individual timing of breeding (relative laying date) and adult survival

To assess the effect of intra-population variation in laying dates, a multistate dataset of capture histories for our population was created. The different states were defined as the quartiles of laying date distribution of the population in a given year, for instance the 25% earliest breeders in the population for a given year were given the state “1” for this year, and the latest quarter were given state “4”. Observations without any laying date information were removed (4 observations, 3 individuals, out of 2258 observations).

The starting model used for model selection (Table 3, Model 85) was set to have only sex allowed as a possible constraint for recapture probabilities (*p*) based on the above selection procedure on the monostate model. It included (1) all possible interactions between state, sex and age for the probability estimates of being in a given initial state (IS); (2) interactions between sex, age, departure and arrival state for between state transition probabilities (T); (3) interactions between state, sex and age for survival probabilities (*φ*). Selection was made sequentially (i.e. by retaining the best structure at each step for the next step) firstly on initial state distribution constraints, then on transition probabilities constraints. For survival constraints, the possible interactive effects of sex and age with the states were tested. Then various potential groupings of states were tested with the association of sex and age as in the best previous models, including grouping of quartiles 1 vs 2 3 4, 1 2 3 vs 4, 1 vs 2 3 vs 4, 1 2 vs 3 4, 1 vs 2 vs 3 4, 1 2 vs 3 vs 4.

#### c Effects of population-level annual covariates on the relative laying date - adult survival link

Based on the model selection described above, we started from a model taking into account differences in laying date quartiles (Tables 3 and 4: model 43), and ran a further model selection by adding four population-level annual metrics (Temp, medLD, popD, repS) and their interactions as continuous temporal covariates (Table 4).

#### d Links between individual breeding success, relative laying date and adult survival

To assess the role of intra-population reproductive investment in the system, another multistate dataset of capture histories was used. The 6 states took into account combinations of the individual differences in reproduction in three categories (Failure: clutch abandoned or predated, 0 fledglings; Low reproduction: number of fledglings below the median number of fledglings per brood in the population this year; High reproduction: number of fledglings above this median) and individual differences in phenology in two categories (Early breeders: first quartile of laying date (state 1 in the previous dataset), Late breeders: second to fourth quartiles of laying date (states 2,3 and 4 in the previous dataset)).

The starting model for the selection included the interactive effects of age, sex and state for its various structural components (Table 5, model 276). Recapture probabilities were assumed to depend on sex only. The sequential selection procedure was the same as above. For survival rates, various groupings of states were tested (Table 5) including grouping of states 1 2 vs 3 4 5 6, 1 2 vs 3 4 vs 5 6, 1 3 5 vs 2 4 6, 1 2 vs 3 5 vs 4 6.

## Results

### Goodness-of-fit

No significant violation of standard models assumptions was found while performing the goodness-of-fit tests using program U-CARE (Choquet et al., 2009a): Detailed goodness-of-fit results are provided in Appendix 1.

#### a Link between population-level phenology and average annual adult survival

The best model from our preliminary selection on the single state capture histories (without covariates) is presented in Table 2A (Model 1). This model assumes an effect of sex only on recapture probabilities, with higher recapture rates for females (*p_males_* = 0.673 [0.611; 0.729]_95%CI_; *p_females_* = 0.859 [0.805; 0.899]_95%CI_), and of age only on survival probabilities, with higher survival probabilities for first year adults than older adults (ϕ_1year_ = 0.597 [0.546; 0.645]_95%CI_; ϕ_2+years_ = 0.526 [0.494; 0.558]_95%CI_).

When adding the three population-level annual demographic covariates (median laying date of the population (medLD), population density (popD) and mean number of fledglings per attempted brood in the population (repS)) as determinants of survival, we observed that adding each covariate separately to Model 1 improved the fit to the data (Table 2B, models 35, 36 and 37). The best model from this selection was Model 23, which took into account the effects of the medLD, popD and their interaction.

All models showed a positive relationship between the median laying date and adult survival and a negative relationship between density and survival, which means that a late reproduction of the population was associated with a high subsequent adult survival, and that a high density of the population was associated with a low subsequent survival (Figure 1A and B). Importantly, these relationships were not due to a change in the proportion of age classes between years and remained unchanged within each age class (see Appendix 3 for additional results within age classes). The interaction between population density and median laying date was negative: when population density increased, the strength of the (positive) relationship between median laying date and adult survival decreased and it almost became negative at very high density.

**Figure 1:**
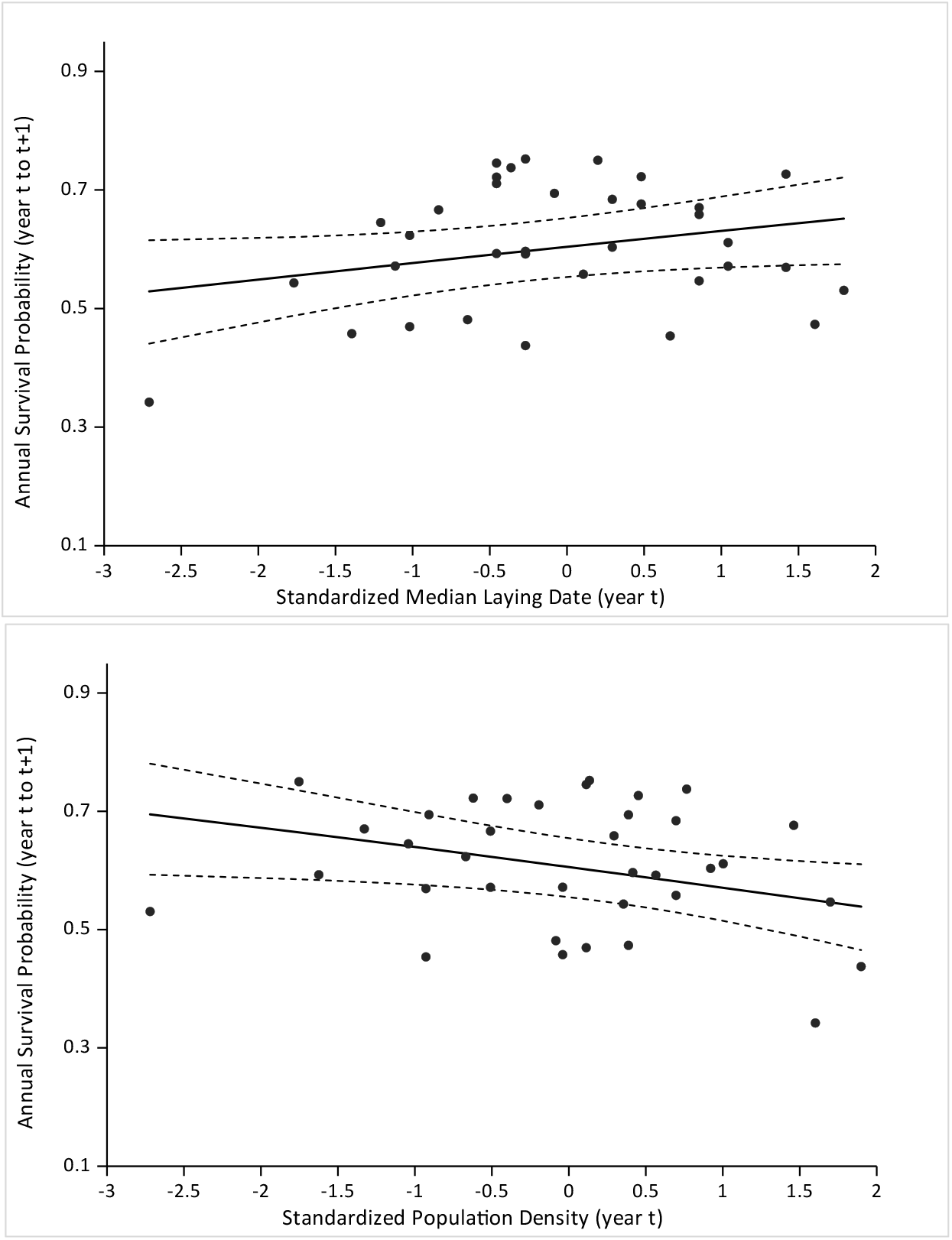
Relationship between annual survival probabilities and two demographic parameters. A: Survival is positively correlated with median laying date of the population in the previous spring (β=0.11; SE=0.06). The points are the estimates of Model 3 (Table 2A) for age 1 with 95% confidence intervals, the solid line is for age 1 from Model 36 (Table 2B) which has a continuous effect of median laying date on survival, its 95% CI are represented by the dashed lines. B: Survival is negatively correlated with population density during the previous spring (β=-0.14; SE=0.07). The points are the estimates of Model 3 (Table 2A) for age 1 with 95% confidence intervals, the solid line is for age 1 from Model 35 (Table 2B) which has a continuous effect of population density on survival, its 95% CI are represented by the dashed lines.

#### b Link between individual timing of breeding (relative laying date) and adult survival

The model selection on the state-separated laying date quartiles capture histories dataset did not show any model stand out in terms of AICc. Seven different models were included within the first 2 points of AICc of our selection (Table 3). All differ only by their constraints on survival estimates, differences among phenological quartiles are included in 5 of these models and the effect of sex is included in only one of them. A model averaging conducted on these models provided an accurate graphic representation of the estimates they produced (Figure 2). While little differences occurred for first year adults, survival for older adults decreased consistently from the first to the last laying date quartile.

**Figure 2:**
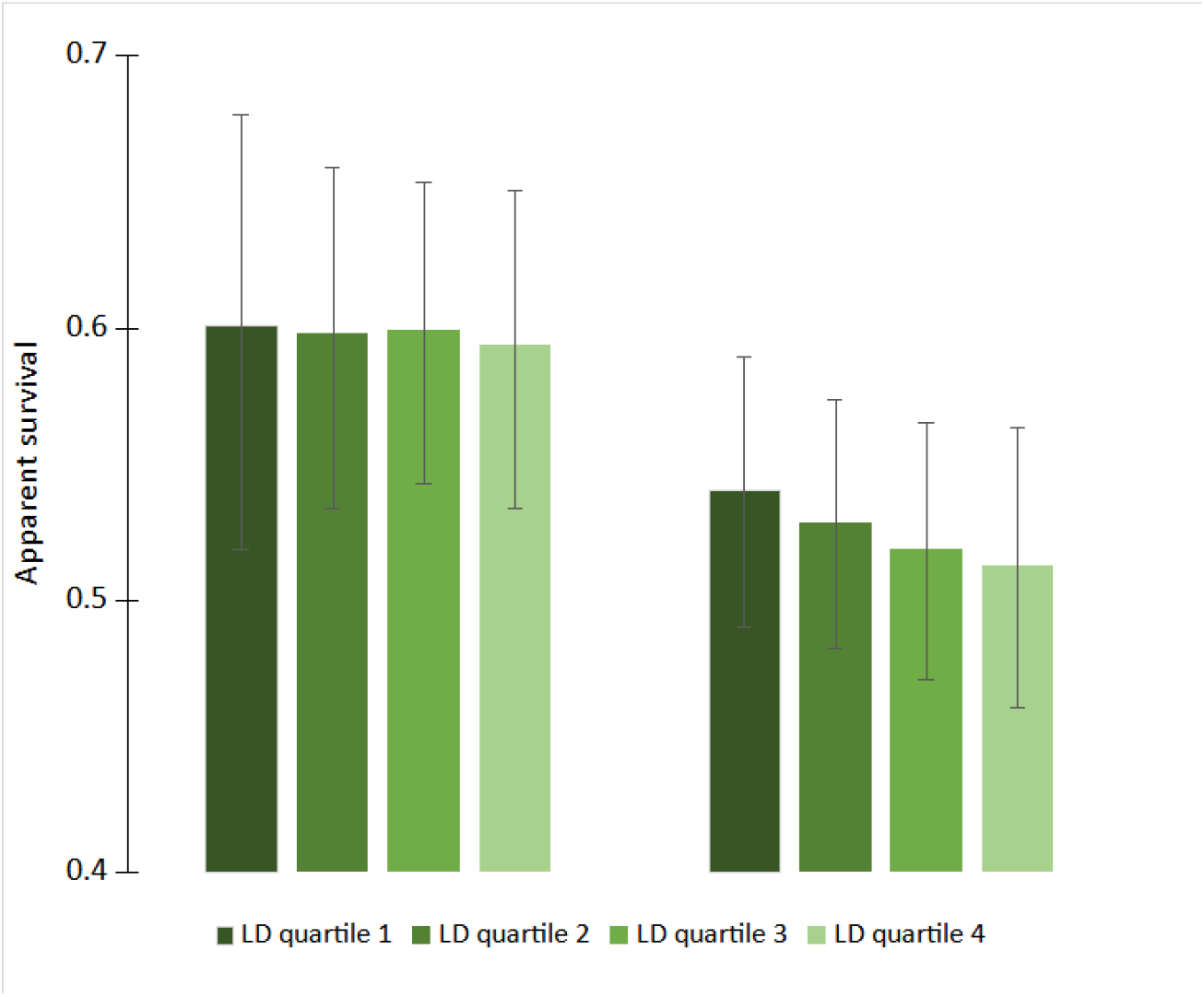
Survival probability estimates associated with different laying date quartiles (LD quartile) of the population obtained by model averaging of the first seven models of our selection (Table 3). For both age classes, each bar represents a different quartile, lines indicate the 95% confidence intervals of the estimates.

#### c Effects of population-level annual covariates on the relative laying date - adult survival link

Performing a new model selection after adding one meteorological (Temp) and three demographic (medLD, popD and repS) continuous temporal covariates to constrain survival probabilities yielded the same results as in the single state model selection (see a)): the most important temporal covariates were the median laying date, population density and their interaction, all included in the best model (Table 4, model 104).

In all models of this selection, the survival estimates associated with laying date quartile 1 (25% earliest breeders of the population for a given year) were higher than those associated with grouped laying date quartiles 2, 3 and 4 (e.g. Model 43 gives ϕ_Age1,LD1_ = 0.628 [0.553; 0.698]_95%CI_; ϕ_Age1,LD234_ = 0.592 [0.541; 0.641]_95%CI_).

Overall, while adult survival was globally positively correlated with median laying date, we observed differences in the slope of this correlation between early breeders and the other individuals: early breeders tended to be almost unaffected by the median laying date (β_LD1_=0.02; SE=0.13 for early breeders (laying date quartile 1) and β_LD234_=0.14; SE=0.07 for others in model 154, Figure 3). The correlation between adult survival and population density remained almost equal for both categories of laying date (β_LD1_=-0.19; SE=0.14 and β_LD234_=-0.13; SE=0.07 in model 139, Table 4).

**Figure 3:**
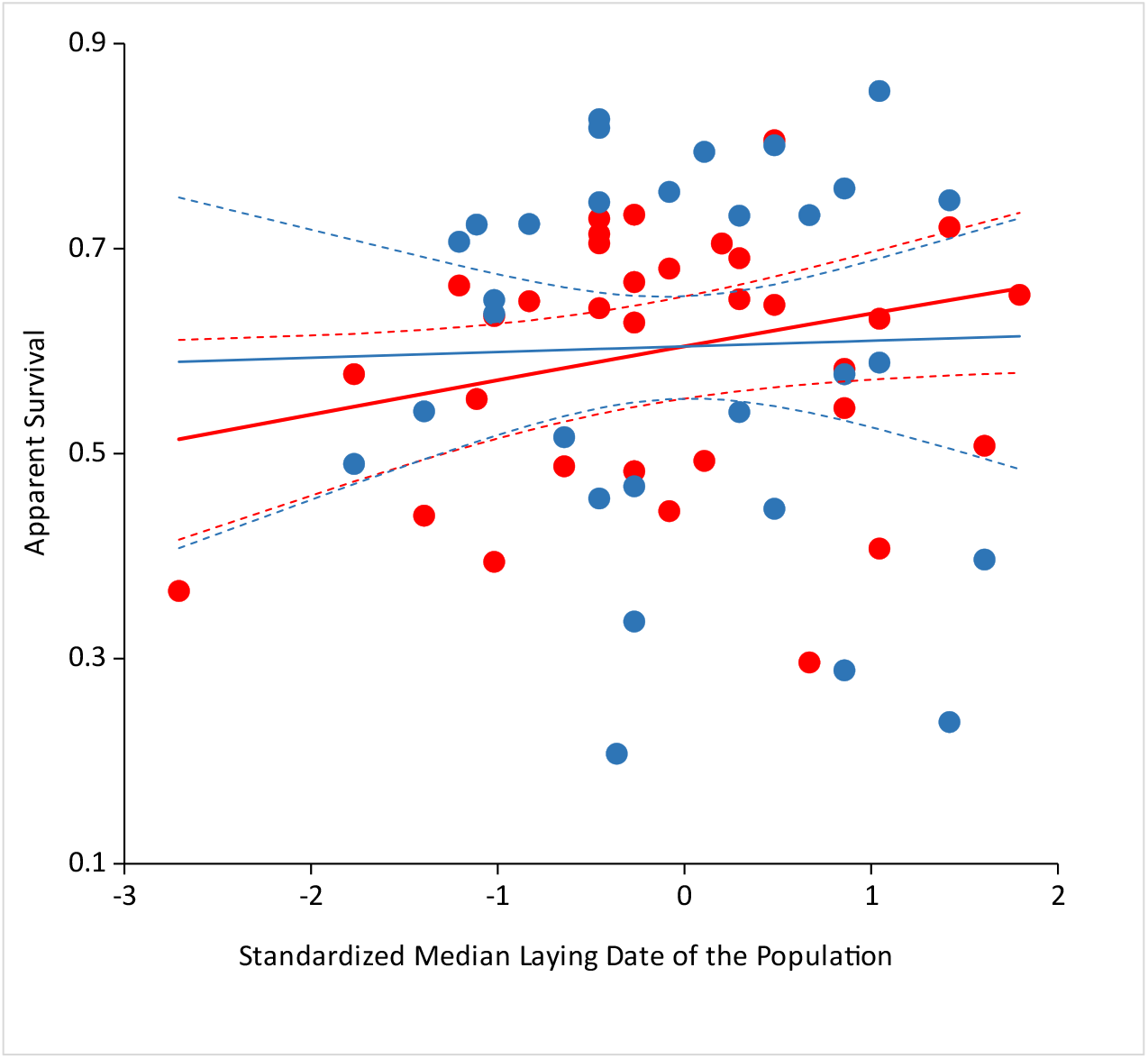
Annual survival probabilities as functions of the median laying date of the population for early breeders (laying date quartile 1, in blue) and for other individuals (laying date quartiles 2, 3 and 4, in red). Filled circles are the annual survival estimates for age 1 from Model 219 (Table 4), solid lines are the regression slopes for age 1 of both laying date quartile groups (1 vs 2,3 and 4) from Model 154 (Table 4), their 95% confidence intervals are represented by the dashed lines.

When looking at the same correlations of annual demographic covariates with survival, but taking the most important covariates namely median laying date, population density and their interaction into account at the same time, different effects arose (Figure 4): For all relative laying date states, at high population density, the positive correlation between survival and the median laying date still occurred. At low population density, the correlation became negative. This pattern was qualitatively similar but stronger for early breeders than for others.

**Figure 4:**
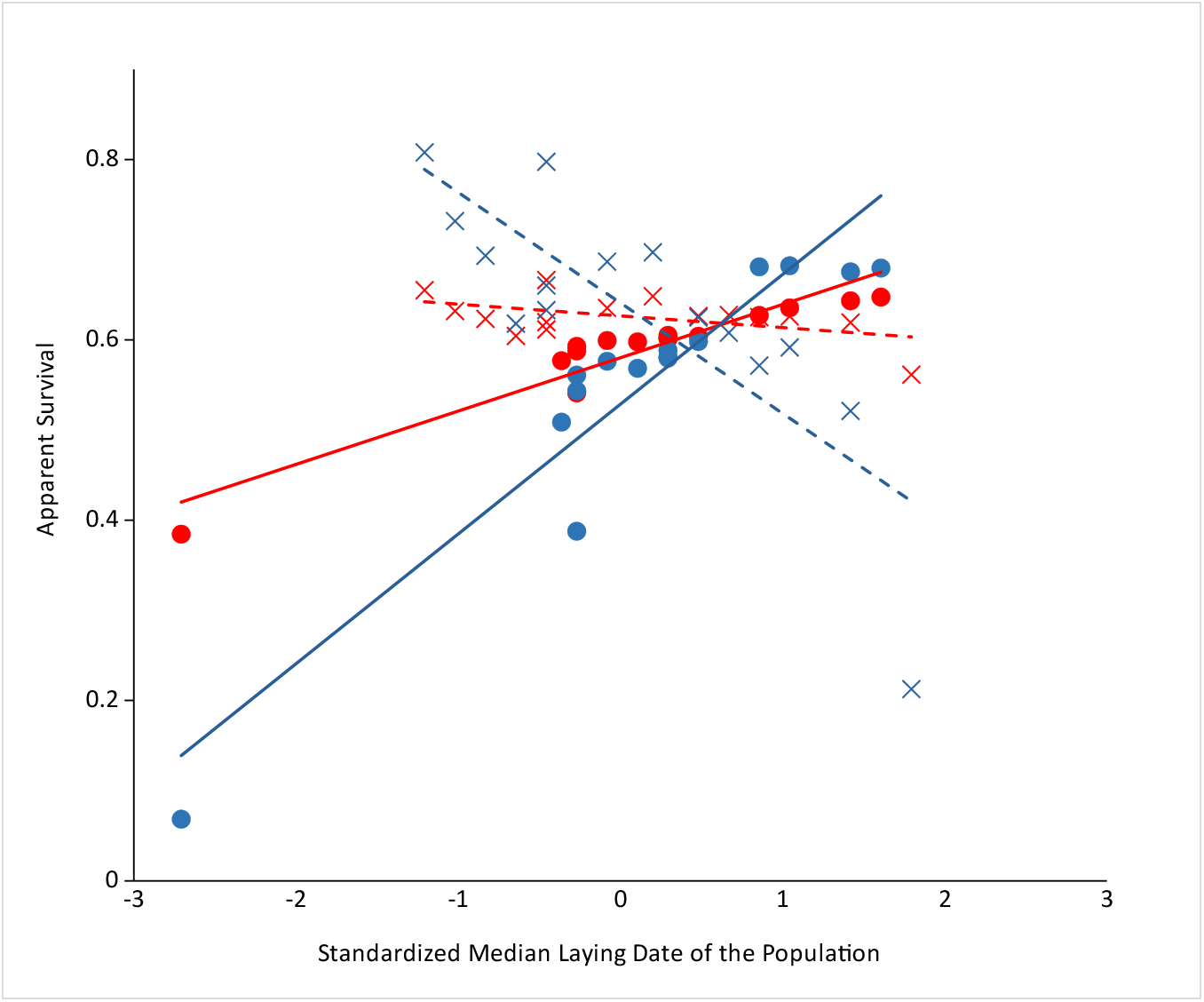
Annual survival probabilities as functions of the median laying date of the population at high (full symbols, solid lines) and low (open symbols, dashed lines) population density, for early breeders (LD quartile 1, blue) and later breeders (LD quartiles 2, 3 and 4, orange). High/low density years are the 15 years of monitoring with highest/lowest population densities. Estimates are from the best model (Model 104, Table 4).

#### d Links between individual breeding success, relative laying date and adult survival

When considering combinations of breeding success and relative laying date states (see Figure 5A for a description of the states) on subsequent survival probability, model selection resulted in four models differing from less than two points of AICc (Table 5, Figure 5C), clearly indicating that the most important survival variation within our combined states occurred between individuals failing to reproduce and those that succeed (regardless of their relative laying date, Figure 5D). The average survival estimates were respectively 0.614 [0.562; 0.664]_95%CI_ for successful breeders and 0.481 [0.387; 0.576]_95%CI_ for failed breeders (estimates from model 220 in Table 5). While early breeders exhibited slightly higher survival probabilities than late breeders for the two categories of successful breeders (Figure 5C and D), our analysis indicated that the magnitude of this difference was small as compared with the difference in survival between failed and successful breeders.

**Figure 5:**
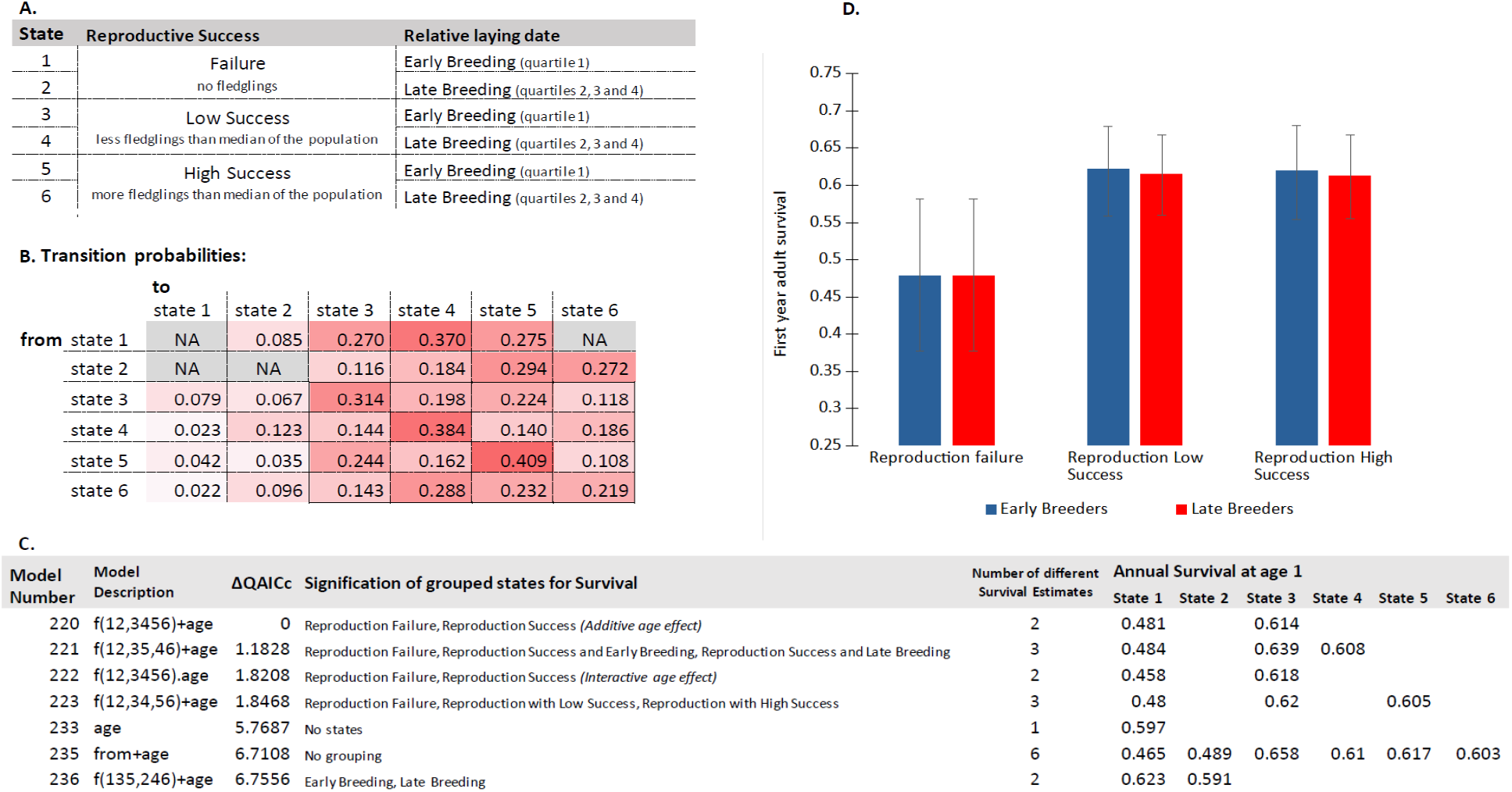
Links between individual breeding success, relative laying date and subsequent adult survival. A: Description of the reproductive success and laying date states used in the multistate models. B: Estimates of transition probabilities among breeding success and relative laying date states (model 220, Table 5 and C). For example, the probability of transitioning from state 3 to state 4 is 0.198. NA correspond to transitions that could not be estimated due to insufficient data. C: Models of significant interest from the model selection (Table 5) with the interpretation of the provided estimates. D: Estimates of first year adult survival in the year following a given breeding success and relative laying date. The provided estimates were obtained by averaging the 4 best models of our selection (Table 5), 95%CIs are represented by the grey lines.

Our analysis also revealed that transitions were asymmetrical with respect to the breeding state: both failed and successful breeders tended to transition preferentially to successful breeding states (model 265, comparison with null model 284 of equal transition probabilities: ΔAICc=138.95, see Figure 5B for transition estimates).

## Discussion

The analysis conducted on the links between reproductive phenology and subsequent adult survival in a Mediterranean forest Blue tit (*Cyanistes caeruleus*) population revealed that (a) at the population scale, early median breeding years are followed by lower average adult survival, (b) at the individual level, earlier breeders within the population have higher subsequent survival than late breeders, (c) population density influences the relative phenology ↔ survival relationship, (d) individual relative reproductive success is also related to subsequent adult survival in our population, in particular, failed reproduction is linked to lower subsequent survival.

### a At the population scale, early breeding years are followed by low average survival

We showed that at the population level, advanced phenology is associated with lower overall subsequent survival in breeders (Table 2, Figure 1A). Although the link between avian breeding phenology and survival has not been examined nearly as much as the link between phenology and reproductive success, this first result is in agreement with previous results in birds (Brinkhof et al., 2002; Nilsson, 1994). Such inter-annual, population-level variation in phenology reflects the average, or common response of breeders in the year t to environmental conditions, including spring temperatures (Bonamour et al., 2019). Similarly, reduced survival between year t and t+1 reflects the average demographic response of breeders to these conditions, regardless of their relative position in the laying date distribution of year t. Reduced average survival may be the consequence of a high energetic cost of early breeding itself, in addition to the cost of high reproduction associated with early breeding (Brinkhof et al., 2002). These results are compatible with our preliminary hypothesis of the importance of a cost of reproduction in our population, as well as with the general prediction that, in short-lived species, costs of reproduction mainly occur on survival (rather than on future reproduction, Hamel et al., 2010; Le Coeur et al., 2018). However, it is important to note that the correlative nature of our analysis prevents us from demonstrating fully this hypothesis. We could be observing, for example, the mortality consequences of a high hydric or energetic stress in warm years, or of high adult predation (due to favourable meteorological conditions for predators) in these years of early avian phenology. Another explanation for the observed pattern could be the existence of phenological mismatches: in particularly warm years, the advance of phenology of the tit population could be insufficient compared to the advance of their prey (Visser and Gienapp, 2019). However, such predator-prey mismatch would also result in a lower reproduction for blue tits, which is not what is generally observed for early years: on the contrary, in the E-Pirio population, the mean number of fledglings per attempted brood increased when the median laying date decreased (+0.12 fledglings/earlier day, *p-value* < 10^-3^).

**Table 2:** Monostate model selection assessing the link between population-level phenology and average annual survival. Full tables are provided in Table A2 of Appendix 2. A: Preliminary model selection assessing the effects of year, sex and age on recapture (*p*) and survival (*φ*) probabilities. Starting model: Model 22, best model: Model 1. AICc(Model 1) = 2550.3171. B: Secondary model selection assessing the effect of three annual population-level demographic covariates (repS= reproductive success, popD = population density, medLD = median laying date) on survival probabilities. The covariates and their interactions were added to preliminary selected model 1. Starting model: Model 1, best model: Model 23. AICc(Model 23) = 2541.4519. np: number of parameters

| Model | Model Description | np | Deviance | $\Delta AICc$ |
| --- | --- | --- | --- | --- |
| **Table 2A** |  |  |  |  |
| **1** | **$\phi$ (age) P(sex)** | **5** | **2540.29** | **0** |
| 2 | $\phi$ (sex + age) P(sex) | 6 | 2540.20 | 1.92 |
| 3 | $\phi$ (age + year) P(sex) | 41 | 2470.48 | 3.57 |
| 4 | $\phi$ (.) P(sex) | 4 | 2545.74 | 3.44 |
| 5 | $\phi$ (sex.age) P(sex) | 7 | 2540.11 | 3.84 |
| 6 | $\phi$ (sex + age + year) P(sex) | 42 | 2470.37 | 5.54 |
| 7 | $\phi$ (sex) P(sex) | 5 | 2545.72 | 5.42 |
| 8 | $\phi$ (sex.age + year) P(sex) | 43 | 2470.18 | 7.41 |
| ... |  |  |  |  |
| 18 | $\phi$(sex.age + sex.year + age.year) P(sex) | 112 | 2407.88 | 92.23 |
| ... |  |  |  |  |
| **22** | **$\phi$(sex.age + sex.year + age.year) P(sex.year)** | **180** | **2305.75** | **143.69** |
| **Table 2B** |  |  |  |  |
| **23** | **$\phi$(age + popD + medLD + popD\*medLD) P(sex)** | **8** | **2525.39** | **0** |
| 24 | $\phi$(age + repS + popD + medLD + popD*medLD) P(sex) | 9 | 2523.60 | 0.22 |
| 25 | $\phi$(age + repS + popD + repS*popD) P(sex) | 8 | 2527.02 | 1.63 |
| 26 | $\phi$(age + repS + popD + medLD + repS*medLD + popD*medLD) P(sex) | 10 | 2523.03 | 1.66 |
| 27 | $\phi$(age + repS + popD + medLD + repS*popD + popD*medLD) P(sex) | 10 | 2523.60 | 2.23 |
| 28 | $\phi$(age + repS + popD + medLD + repS*popD) P(sex) | 9 | 2526.36 | 2.98 |
| 29 | $\phi$(age + repS + popD + LEDmed + repS*popD + repS*medLD + popD*medLD) P(sex) | 11 | 2523.01 | 3.66 |
| ... |  |  |  |  |
| 35 | $\phi$(age + popD) P(sex) | 6 | 2535.62 | 6.19 |
| 36 | $\phi$(age + medLD) P(sex) | 6 | 2536.86 | 7.44 |
| 37 | $\phi$(age + repS) P(sex) | 6 | 2537.07 | 7.65 |
| ... |  |  |  |  |
| **1** | **$\phi$(age)P(sex)** | **5** | **2540.29** | **8.86** |

### b At the individual level, relative early breeders have higher survival the following year

Interestingly, when examining the relationship between survival and relative laying date (i.e., the position of the focal individual within the laying date distribution in a given year), the opposite result was obtained compared to the population scale. Regardless of the median laying date of the population, individuals that breed early in the population have higher subsequent survival than others (for ages of two years or more) (Figure 2). This result suggests that some individual heterogeneity (Gimenez et al., 2018) occurs within our study population: while breeding early can be seen as a strategy to improve breeding success (Marrot et al., 2018) that may convey some survival cost (as suggested by our present population-level results), early breeders may be more prone to survive in spite of this cost. Similar results have been obtained by previous studies on the cost of reproduction: while reproduction is expected to be costly, positive covariation between breeding success and subsequent survival and other ‘big house big car’ correlations are frequently observed in free ranging animal populations (van Noordwijk and de Jong, 1986; Reznick et al., 2000; Robert et al., 2015). In our blue tit population, the high repeatability of the rank of laying date among years (R = 0.406 [0.316; 0.493]_95%CI_ for females, see Appendix 4 for details) suggests indeed the occurrence of some intrapopulation heterogeneity regarding the phenological strategy. This result is confirmed by the analysis of the transitions among states of relative laying date, which are not random (comparison of model 74 and model 101, ΔAICc = 113.33, Table 3). In particular, the earliest breeders (quartile 1 of the laying date distribution at year t) have a very high probability (52%) to remain in quartile 1 at year t+1 (see transition probabilities matrix in Appendix 5). These results are in line with previous findings in many bird species that breeding phenology is repeatable and heritable (see e.g. Postma, 2014; for the focal population Delahaie et al., 2017 estimated a heritability of laying date of 0.125 [0.059; 0.199]_95%CI_).

**Table 3:** Multistate model selection assessing the link between relative laying date and subsequent adult survival. Full table is provided in Table A3 of Appendix 2. The four states represent laying date quartiles (an individual in state 1 has laid in the earliest 25% of the population). Starting model: Model 85, best model: Model 41. AICc(Model 41) = 8586.2435. For initial state distribution: “to” means that at their first capture, individuals are not evenly distributed among the four phenological quartiles. For between states transition probabilities: “f.to” means that transition probabilities depend on the departure state (f), the arrival state (to) and their interaction. For survival: “f” means that survival depends on the current state of the individual. np: number of parameters.

| Model | Model Description | np | Deviance | $\Delta AICc$ |
| --- | --- | --- | --- | --- |
| 41 | IS(to.age) $\phi$(age) T(f.to + age) P(sex) | 23 | 8539.83 | 0 |
| 42 | IS(to.age) $\phi$(f(1 2 3,4) + age) T(f.to + age) P(sex) | 24 | 8538.58 | 0.79 |
| 43 | IS(to.age) $\phi$(f(1,2 3 4) + age) T(f.to + age) P(sex) | 24 | 8538.59 | 0.80 |
| 44 | IS(to.age) $\phi$(f(1 2,3 4) + age) T(f.to + age) P(sex) | 24 | 8539.17 | 1.38 |
| 45 | IS(to.age) $\phi$(f(1,2 3 4).age) T(f.to + age) P(sex) | 25 | 8537.51 | 1.75 |
| 46 | IS(to.age) $\phi$(sex + age) T(f.to + age) P(sex) | 24 | 8539.77 | 1.98 |
| 47 | IS(to.age) $\phi$(f(1 4,2 3) + age) T(f.to + age) P(sex) | 24 | 8539.83 | 2.04 |
| 48 | IS(to.age) $\phi$(f(1,2 3,4) + age) T(f.to + age) P(sex) | 25 | 8538.02 | 2.26 |
| 49 | IS(to.age) $\phi$(f(1,2 3 4) + sex + age) T(f.to + age) P(sex) | 25 | 8538.54 | 2.79 |
| ... |  |  |  |  |
| 74 | IS(to.age) $\phi$(f.age.sex) T(f.to + age) P(sex) | 37 | 8528.26 | 17.08 |
| ... |  |  |  |  |
| 84 | IS(to.age) $\phi$(f.age.sex) T(f.to.age.sex) P(sex) | 72 | 8491.61 | 53.38 |
| 85 | IS(to.sex.age) $\phi$(f.age.sex) T(f.to.age.sex) P(sex) | 78 | 8483.63 | 58.10 |
| ... |  |  |  |  |
| 101 | IS(to.age) $\phi$(f.age.sex) T(.) P(sex) | 25 | 8666.16 | 130.41 |
| ... |  |  |  |  |

**Table 4:**
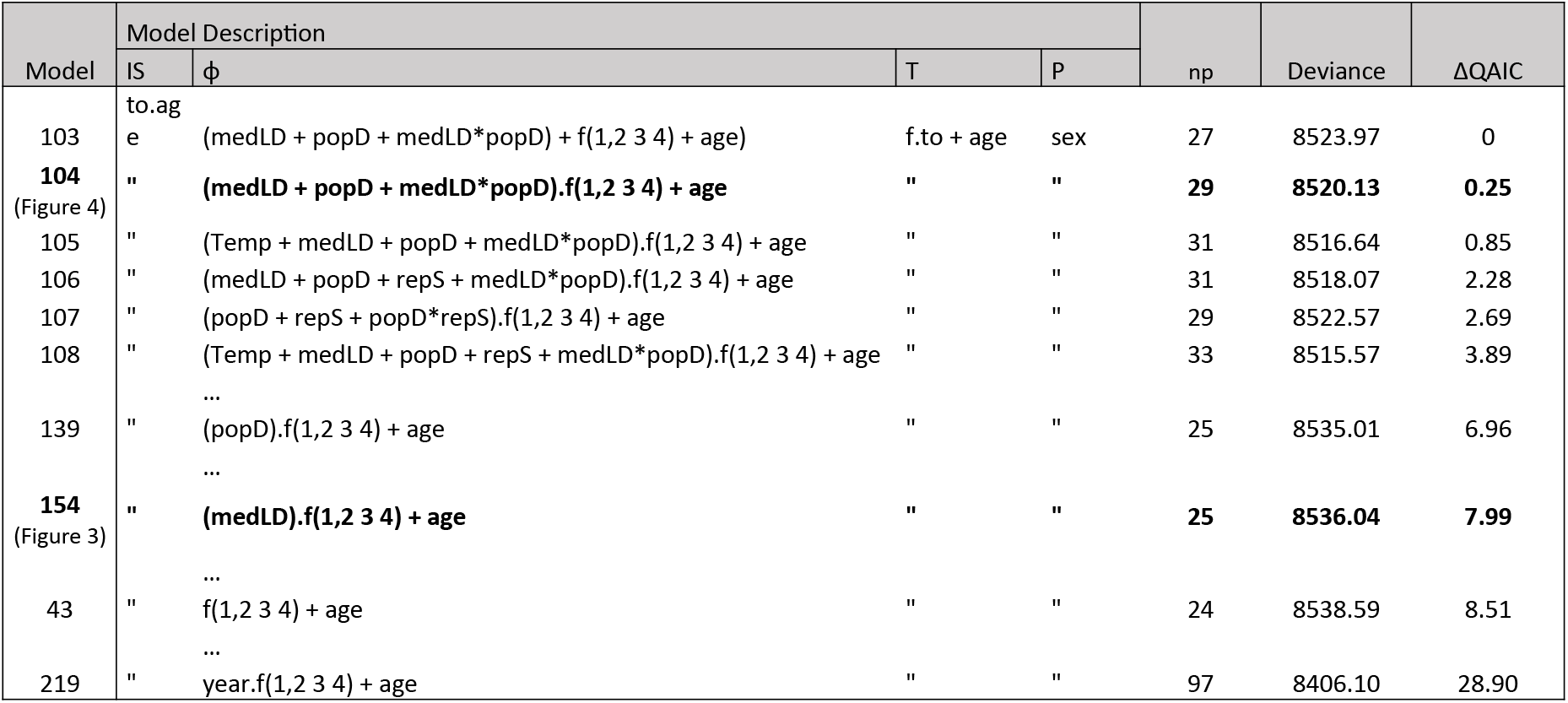
Multistate model selection assessing the effects of population-level annual covariates on the relative laying date - adult survival link. Full table is provided in Table A4 of Appendix 2. The effects of 4 annual covariates (Temp = mean daily mean temperature from March 31^st^ to May 7^th^, repS = reproductive success, popD = population density, medLD = median laying date) and their first-order interactions as continuous temporal covariates on survival probabilities were tested on one of the best models from the previous selection (Model 43, Table 3). Starting model: Model 43, best model: Model 103. AICc(Model 103) = 8578.5326. For initial state distribution: “to” means that at their first capture, individuals are not evenly distributed among the four phenological quartiles.. For between states transitions probabilities: “f.to” means that transition probabilities depend on the departure state (f), the arrival state (to) and their interaction. For survival: “f” means that survival depends on the current state of the individual. np: number of parameters.

**Table 5:** Multistate model selection assessing the links between individual breeding success, relative laying date and adult survival. Full table is provided in Table A5 of Appendix 2. The 6 states represent different combinations of individual breeding success and relative laying date (see Figure 5 for details). Starting model: Model 276 included the interactive effects of age, sex and state for its various structural components. Recapture probabilities were assumed to depend on sex only. Sequential selection procedure was the same as in previous model selection (Table 4). Best model: Model 220. AICc(Model 220) = 9599.8097. np: number of parameters.

| Model | Model Description | np | Deviance | $\Delta AICc$ |
| --- | --- | --- | --- | --- |
| **220** | ***IS(to.age) $\phi(f(12,3456) + age) T(f.to) P(sex)$*** | **45** | **9508.25** | **0** |
| **221** | ***IS(to.age) $\phi(f(12,35,46) + age) T(f.to) P(sex)$*** | **46** | **9507.36** | **1.18** |
| **222** | ***IS(to.age) $\phi(f(12,3456).age) T(f.to) P(sex)$*** | **46** | **9508.00** | **1.82** |
| **223** | ***IS(to.age) $\phi(f(12,34,56) + age) T(f.to) P(sex)$*** | **46** | **9508.03** | **1.85** |
| 224 | IS(to.age) $\phi(f(12,3456) + sex + age) T(f.to) P(sex)$ | 46 | 9508.19 | 2.01 |
| 225 | IS(to.age) $\phi(f(12,3456).age + sex) T(f.to) P(sex)$ | 47 | 9507.96 | 3.84 |
| 226 | IS(to.age) $\phi(f(12,34,56) + sex + age) T(f.to) P(sex)$ | 47 | 9507.98 | 3.86 |
| 227 | IS(to.age) $\phi(f(12,3456).sex + age) T(f.to) P(sex)$ | 47 | 9507.98 | 3.87 |
| 228 | IS(to.age) $\phi(f(12,3456) + sex.age) T(f.to) P(sex)$ | 47 | 9508.16 | 4.05 |
| 229 | IS(to.age) $\phi(f(12,3456)) T(f.to) P(sex)$ | 44 | 9514.23 | 3.91 |
| ... |  |  |  |  |
| **233** | ***IS(to.age) $\phi(age) T(f.to) P(sex)$*** | **44** | **9516.09** | **5.77** |
| ... |  |  |  |  |
| **235** | ***IS(to.age) $\phi(from + age) T(f.to) P(sex)$*** | **49** | **9506.67** | **6.71** |
| **236** | ***IS(to.age) $\phi(f(135,246) + age) T(f.to) P(sex)$*** | **45** | **9515.01** | **6.76** |
| ... |  |  |  |  |
| 265 | IS(to.age) $\phi(f.age.sex) T(f.to) P(sex)$ | 66 | 9498.25 | 33.79 |
| ... |  |  |  |  |
| 274 | IS(to.age) $\phi(f.age.sex) T(f.to.age.sex) P(sex)$ | 153 | 9414.88 | 139.55 |
| ... |  |  |  |  |
| 276 | IS(to.sex.age) $\phi(f.age.sex) T(f.to.age.sex) P(sex)$ | 163 | 9408.55 | 155.79 |
| ... |  |  |  |  |
| 284 | IS(to.age) $\phi(f.age.sex) T(.) P(sex)$ | 37 | 9697.50 | 172.74 |
| ... |  |  |  |  |

When considering both the absolute (median of the population) and the relative laying date, we found that the strength of the correlation between survival and the median laying date varies between early breeders and later breeders (Figure 3). Early breeders, on top of having higher survival rates, are less affected by the “quality” of the year (warm or cold spring resulting in early or late median laying date) than late breeders. This result confirms that earlier breeders might be high resource individuals able to secure the earliest and best territories while also having high somatic maintenance.

### c Population density influences the relative phenology ↔ survival relationship

Population density was another important factor (negatively) correlated with annual subsequent survival (Figure 1B), presumably reflecting the intensity of intraspecific competition for food resources (Gamelon et al., 2016). We thus included density in the multistate individual model to test whether population breeding density modulated the relationship between phenology and survival probability. Results show that at high breeding population density the suspected potential trade-off between early reproduction and subsequent survival becomes apparent (Figure 4): this suggests that the unfavourable environment of high intra-population competition increases the costs of early breeding. On the contrary, at low population density, the constraints of the trade-off seem to be relaxed: no cost in survival of breeding early can be observed, and in warm years (early median laying date), on top of having better reproduction (Delahaie et al., 2017), early breeding individuals have higher survival rates (Figure 4).

Two main conclusions can be drawn from these results. First, the cost of breeding early is modulated by environmental conditions. Although, to our knowledge, it has never been applied to phenology, this finding is in agreement with previous results showing that costs of reproduction may be modulated by age (Rauset et al., 2015), resource availability (Le Coeur et al., 2018) or meteorological conditions (Stoelting et al., 2014). Second, while the cost of breeding earlier than the other breeders in a given year may be hidden by demographic heterogeneity under average conditions (Figure 2), it becomes apparent under very harsh conditions (here, a combination of warm spring and high population density, which are both correlated to low adult survival). Under such harsh conditions, individuals that invest in early breeding show particularly low survival probabilities (Figure 4). Again, similar interplays between demographic heterogeneity, reproduction cost and environmental conditions have been found in previous studies (Robert et al., 2012). For example, in the Alpine ibex (Capra ibex), survival probabilities were found higher for successfully breeding females than for unsuccessful females under average conditions. However, during a severe disease outbreak, survival patterns according to reproductive state were reversed, with lower survival for successful than unsuccessful females (Garnier et al., 2016).

### d Both individual laying date and individual breeding success are related to survival

Life-history theory predicts that, at the scale of organisms, investment in current reproduction may constrain the energy available for maintenance, leading to trade-offs with survival and/or future reproduction (Roff, 1993; Stearns, 1992) referred to as the cost of reproduction. In the context of our blue tit population, if early breeding is a strategy for breeders to improve their reproductive success, the survival cost associated with early reproduction can be considered a form of reproductive cost. However, it is notoriously difficult to disentangle the cost of early reproduction *per se* from the cost associated with high reproductive success. Previous analyses in our study population indicate that early breeders are generally more successful than late breeders (Porlier et al., 2012), which implies that the effects of breeding success and laying date on survival are partially confounded. When considering combinations of reproductive success and relative laying date in the survival models, results uncover strong differences in survival probabilities between individuals with reproduction success (at least one fledgling) or failure (Figure 5). All individuals of our datasets were captured in the nest boxes during reproduction: they have all at least attempted reproduction and reached the nestling stage, i.e. successfully hatched at least one nestling. This means that individuals with failed reproduction (0 fledglings) encountered strong impediments during the breeding event (scarceness of resources, harsh climatic events, predation of the brood, abandonment by one or the other parent, or death).

A high proportion of these brood failures may be linked to the death of one or both parents: as it occurs after the recapture and identification of the parents (otherwise they would not be in the dataset), it will be analysed as death during the subsequent year and therefore we expected to observe low survival for adults with failed reproduction compared to others. However, the positive effect of being an early breeder is still observed here, on top of the effects of reproductive success. Model selection indicates that (1) models assuming survival differences between early and late breeders are part of the set of best models (ΔAICc<2, Table 5) (2) among breeders that fledged at least one chick, the relative laying date explains more deviance than the breeding success (model 221 vs. 223, Table 5).

### e Evolutionary perspectives in a context of climate change

Previous selection analyses using reproductive success as fitness estimator concluded that early breeding individuals are favoured by selection (e.g. Klomp, 1970; Marrot et al., 2018 in the focal population). As laying date is a heritable trait (Delahaie et al., 2017; Postma, 2014), and as it stands under negative selection, many studies using reproductive success were surprised by the absence of evolution of laying date in the context of climate change (e.g. Price et al., 1988). An often suggested explanation to this evolutionary stasis is the potential presence of a positive selection on laying date via survival, opposed to and compensating the negative selection via reproductive success (Merilä et al., 2001). Here, we showed that this cannot be the case in our population, as early breeding individuals have not only higher reproductive success, but also higher survival, which only increases the selection for early breeding.

Another explanation behind common evolutionary stasis in heritable selected traits is the spatio-temporal fluctuations of selection (e.g. Miehls et al., 2015). In our study population, the only case in which selection on survival was inversed was in harsh condition years (early median laying date and high population density), whereby later laying individuals had higher survival rates: If the frequency of such years were to increase in the context of climate change, the selection regime of the focal population might face significant changes.

## Supporting information

Data description file

model files

model description file

## Acknowledgements

We are grateful to Emmanuelle Cam, Aurélie Cohas, Olivier Gimenez, François Chiron, François Sarrazin, Roger Pradel and Pierre-Yves Henry for their constructive comments on an earlier version of this manuscript. We thank all the people who helped conduct the blue tit monitoring in the past four decades, in particular Jacques Blondel, Phillippe Perret, Marcel Lambrechts, Claire Doutrelant, Denis Réale, Samuel Caro, and Christophe de Franceschi (who also managed the database). We thank the APEEM, the ONF and Achille Sanroma for field logistics, and the Fango valley MAB reserve. The long term project has been supported by the OSU-OREME since 2009.

## Appendix 1: Goodness-of-fit

No significant violation of standard models assumptions was found while performing the goodness-of-fit tests using program U-CARE (Choquet et al., 2009): for the monostate dataset χ^2^(df=184)=85.34, P-level=1; for the multistate dataset with relative laying date states: χ^2^(df=98)=21.289 for group 1, χ^2^(df=131)=26.885 for group 2, χ^2^(df=57)=14.276 for group 3, χ^2^(df=36)=2.808 for group 4, p-value=1 for all groups; for the multistate dataset with breeding success and relative laying date states: χ^2^(df=83)=21.318 for group 1, χ^2^(df=132)=36.779 for group 2, χ^2^(df=51)=13.323 for group 3, χ^2^(df=30)=4.530 for group 4, p-value=1 for all groups.

## Appendix 2: Full model selection tables (Tables A2–A5)

**Table A2 A.**

| Model | Model Description | np | Deviance | $\Delta AICc$ |
| --- | --- | --- | --- | --- |
| **1** | **$\phi$ (age) P(sex)** | **5** | **2540.29** | **0** |
| 2 | $\phi$ (sex + age) P(sex) | 6 | 2540.20 | 1.92 |
| 3 | $\phi$ (age + year) P(sex) | 41 | 2470.48 | 3.57 |
| 4 | $\phi$ (.) P(sex) | 4 | 2545.74 | 3.44 |
| 5 | $\phi$ (sex.age) P(sex) | 7 | 2540.11 | 3.84 |
| 6 | $\phi$ (sex + age + year) P(sex) | 42 | 2470.37 | 5.54 |
| 7 | $\phi$ (sex) P(sex) | 5 | 2545.72 | 5.43 |
| 8 | $\phi$ (sex.age + year) P(sex) | 43 | 2470.18 | 7.41 |
| 9 | $\phi$ (year) P(sex) | 40 | 2476.49 | 7.52 |
| 10 | $\phi$ (sex + year) P(sex) | 41 | 2476.47 | 9.56 |
| 11 | $\phi$ (age.year) P(sex) | 74 | 2440.68 | 42.97 |
| 12 | $\phi$ (age.year + sex) P(sex) | 75 | 2440.63 | 45.04 |
| 13 | $\phi$ (sex.age + age.year) P(sex) | 76 | 2440.60 | 47.14 |
| 14 | $\phi$ (sex.year + age) P(sex) | 78 | 2439.65 | 50.45 |
| 15 | $\phi$ (sex.age + sex.year) P(sex) | 79 | 2439.22 | 52.15 |
| 16 | $\phi$ (sex.year) P(sex) | 77 | 2444.39 | 53.06 |
| 17 | $\phi$ (sex.year + age.year) P(sex) | 111 | 2408.41 | 90.56 |
| **18** | **$\phi$(sex.age + sex.year + age.year) P(sex)** | **112** | **2407.88** | **92.23** |
| 19 | $\phi$(sex.age + sex.year + age.year) P(sex + year) | 147 | 2337.96 | 100.25 |
| 20 | $\phi$(sex.age + sex.year + age.year) P(.) | 111 | 2427.16 | 109.31 |
| 21 | $\phi$(sex.age + sex.year + age.year) P(year) | 146 | 2361.01 | 121.03 |
| **22** | **$\phi$(sex.age + sex.year + age.year) P(sex.year)** | **180** | **2305.75** | **143.69** |

**Table A2 B.** Monostate model selection assessing the link between population-level phenology and average annual survival. **A:** Preliminary model selection assessing the effects of year, sex and age on recapture (*p*) and survival (*φ*) probabilities. Starting model: Model 22, best model: Model 1. AICc(Model 1) = 2550.3171. **B:** Secondary model selection assessing the effect of three annual population-level demographic covariates (repS=reproductive success, popD = population density, medLD = median laying date) on survival probabilities. The covariates and their interactions were added to preliminary selected model 1. Starting model: Model 1, best model: Model 23. AICc(Model 23) = 2541.4519. np: number of parameters.

| Model | Model Description | np | Deviance | $\Delta\text{AICc}$ |
| --- | --- | --- | --- | --- |
| **23** | **$\phi(\text{age} + \text{popD} + \text{medLD} + \text{popD} * \text{medLD}) P(\text{sex})$** | **8** | **2525.39** | **0** |
| 24 | $\phi(\text{age} + \text{repS} + \text{popD} + \text{medLD} + \text{popD} * \text{medLD}) P(\text{sex})$ | 9 | 2523.60 | 0.22 |
| 25 | $\phi(\text{age} + \text{repS} + \text{popD} + \text{repS} * \text{popD}) P(\text{sex})$ | 8 | 2527.02 | 1.63 |
| 26 | $\phi(\text{age} + \text{repS} + \text{popD} + \text{medLD} + \text{repS} * \text{medLD} + \text{popD} * \text{medLD}) P(\text{sex})$ | 10 | 2523.03 | 1.67 |
| 27 | $\phi(\text{age} + \text{repS} + \text{popD} + \text{medLD} + \text{repS} * \text{popD} + \text{popD} * \text{medLD}) P(\text{sex})$ | 10 | 2523.60 | 2.24 |
| 28 | $\phi(\text{age} + \text{repS} + \text{popD} + \text{medLD} + \text{repS} * \text{popD}) P(\text{sex})$ | 9 | 2526.36 | 2.98 |
| 29 | $\phi(\text{age} + \text{repS} + \text{popD} + \text{LEDmed} + \text{repS} * \text{popD} + \text{repS} * \text{medLD} + \text{popD} * \text{medLD}) P(\text{sex})$ | 11 | 2523.01 | 3.67 |
| 30 | $\phi(\text{age} + \text{popD} + \text{medLD}) P(\text{sex})$ | 7 | 2531.07 | 3.66 |
| 31 | $\phi(\text{age} + \text{repS} + \text{popD}) P(\text{sex})$ | 7 | 2531.11 | 3.70 |
| 32 | $\phi(\text{age} + \text{repS} + \text{popD} + \text{medLD}) P(\text{sex})$ | 8 | 2529.77 | 4.38 |
| 33 | $\phi(\text{age} + \text{repS} + \text{popD} + \text{medLD} + \text{repS} * \text{popD} + \text{repS} * \text{medLD}) P(\text{sex})$ | 10 | 2526.35 | 4.98 |
| 34 | $\phi(\text{age} + \text{repS} + \text{popD} + \text{medLD} + \text{repS} * \text{medLD}) P(\text{sex})$ | 9 | 2529.57 | 6.19 |
| **35** | **$\phi(\text{age} + \text{popD}) P(\text{sex})$** | **6** | **2535.62** | **6.20** |
| **36** | **$\phi(\text{age} + \text{medLD}) P(\text{sex})$** | **6** | **2536.86** | **7.44** |
| 37 | $\phi(\text{age} + \text{repS}) P(\text{sex})$ | 6 | 2537.07 | 7.65 |
| 38 | $\phi(\text{age} + \text{repS} + \text{medLD}) P(\text{sex})$ | 7 | 2536.02 | 8.61 |
| **1** | **$\phi(\text{age}) P(\text{sex})$** | **5** | **2540.29** | **8.87** |
| 40 | $\phi(\text{age} + \text{repS} + \text{medLD} + \text{repS} * \text{medLD}) P(\text{sex})$ | 8 | 2535.45 | 10.06 |

**Table A3:**
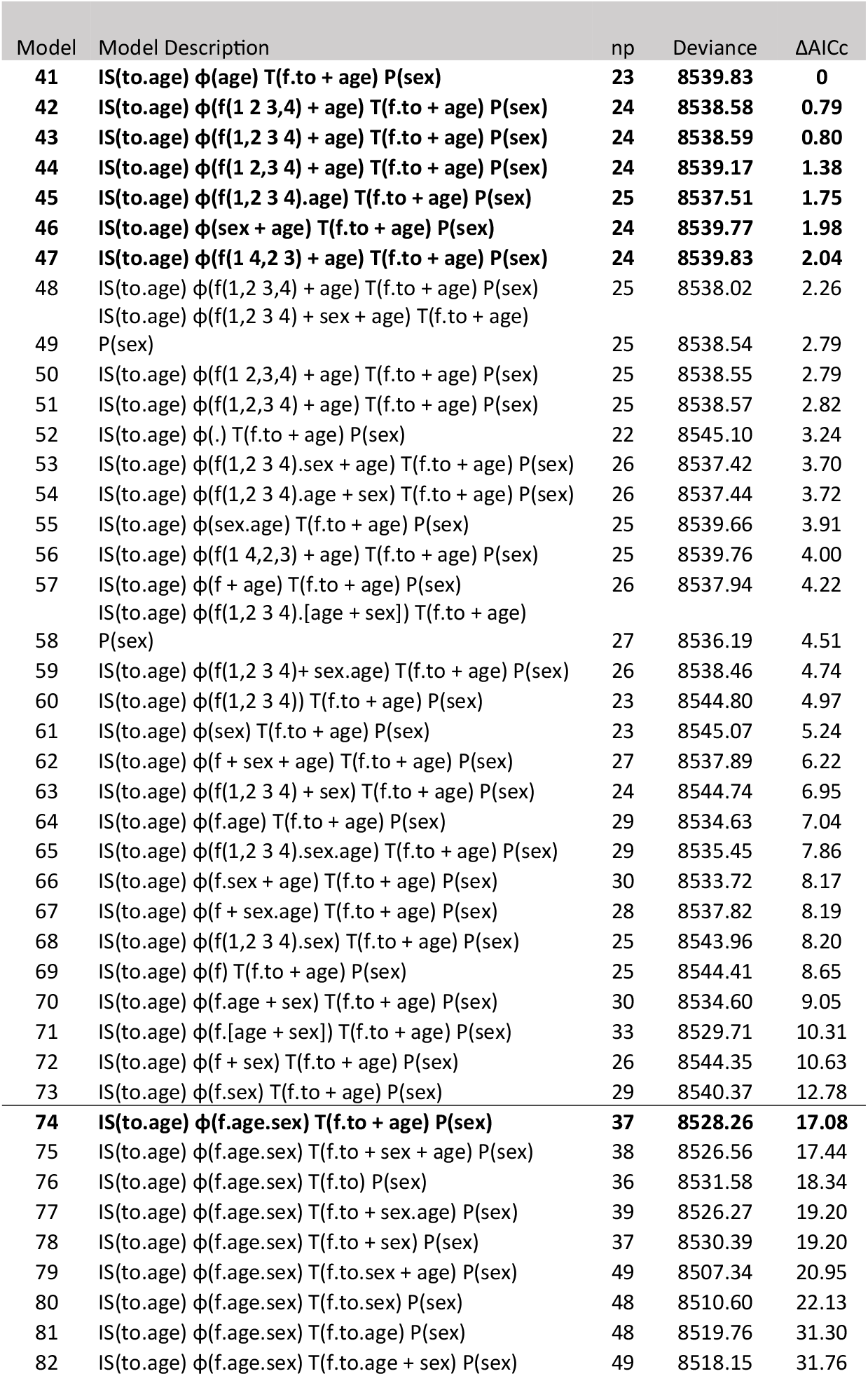
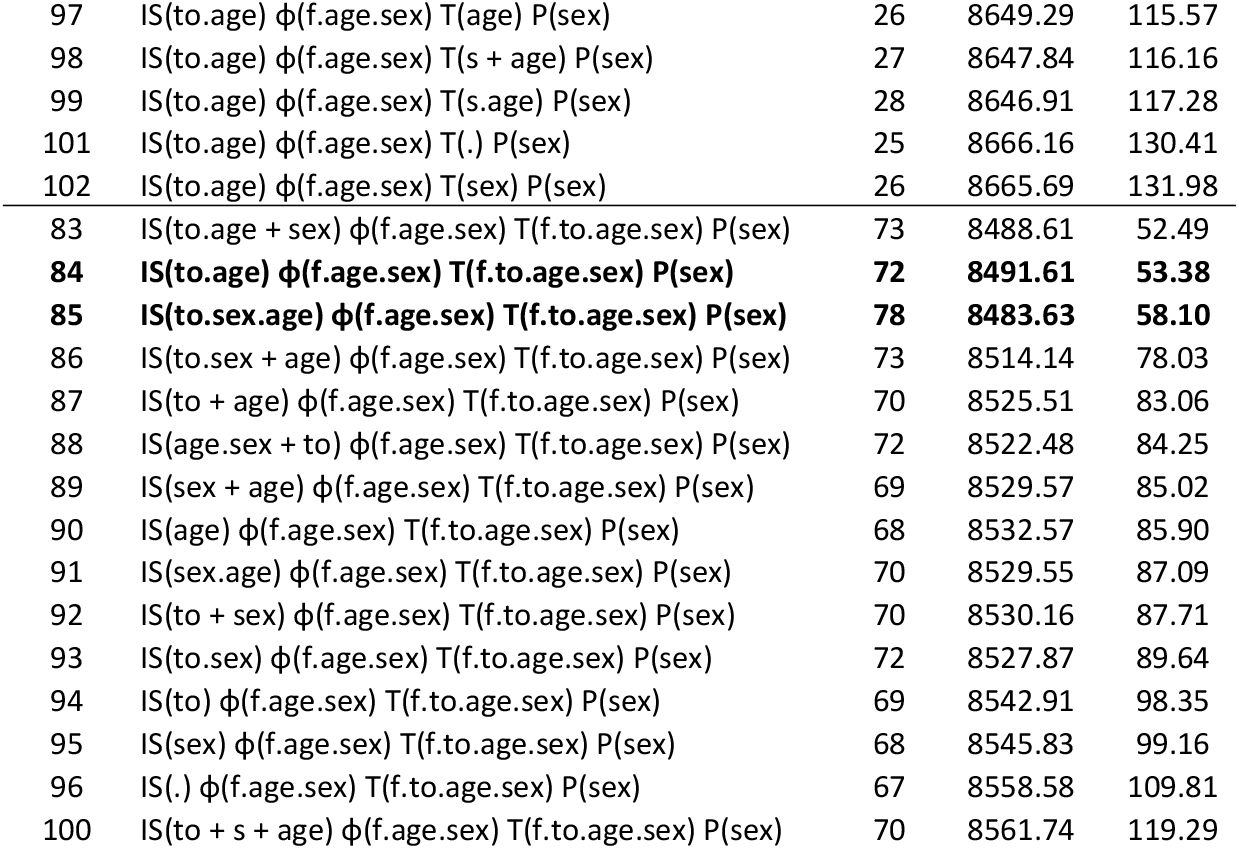
Multistate model selection assessing the link between relative laying date and subsequent adult survival. The four states represent laying date quartiles (an individual in state 1 in year t has laid in the earliest 25% of the population that year). All models have sex as only constraint on recapture probabilities (P). Selection was made sequentially (by retaining the best structure at each step for the next step), firstly on initial state distribution constraints (IS), then on transition probabilities constraints (T), then on survival constraints (ϕ). Starting model: Model 85, best model: Model 41. AICc(Model 41) = 8586.2435. For initial state distribution: “to” means that at their first capture, individuals are not evenly distributed among the four phenological quartiles. For between states transition probabilities: “f.to” means that transition probabilities depend on the departure state (f), the arrival state (to) and their interaction. For survival: “f” means that survival depends on the current state of the individual. np: number of parameters.

**Table A4:**
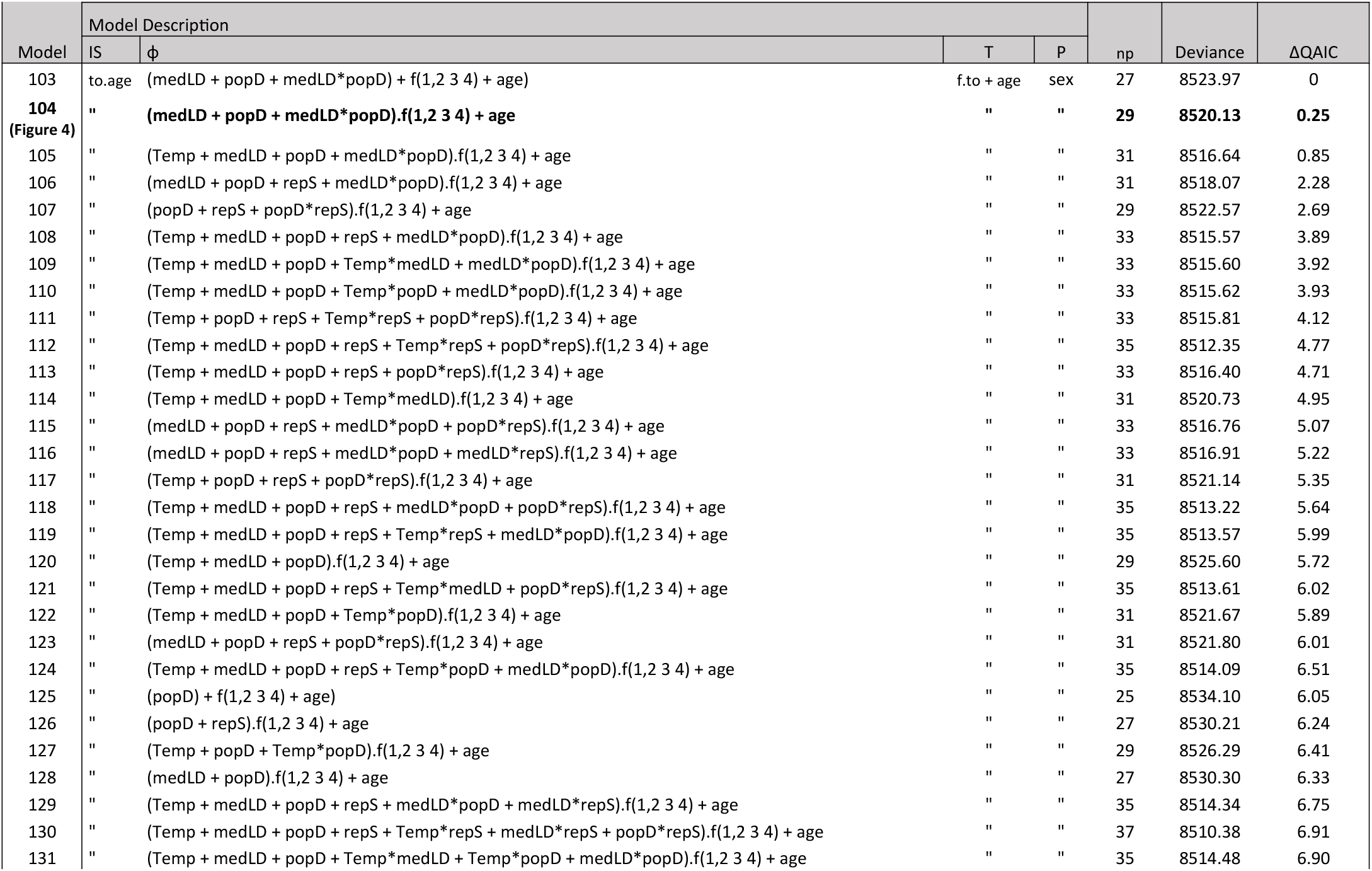
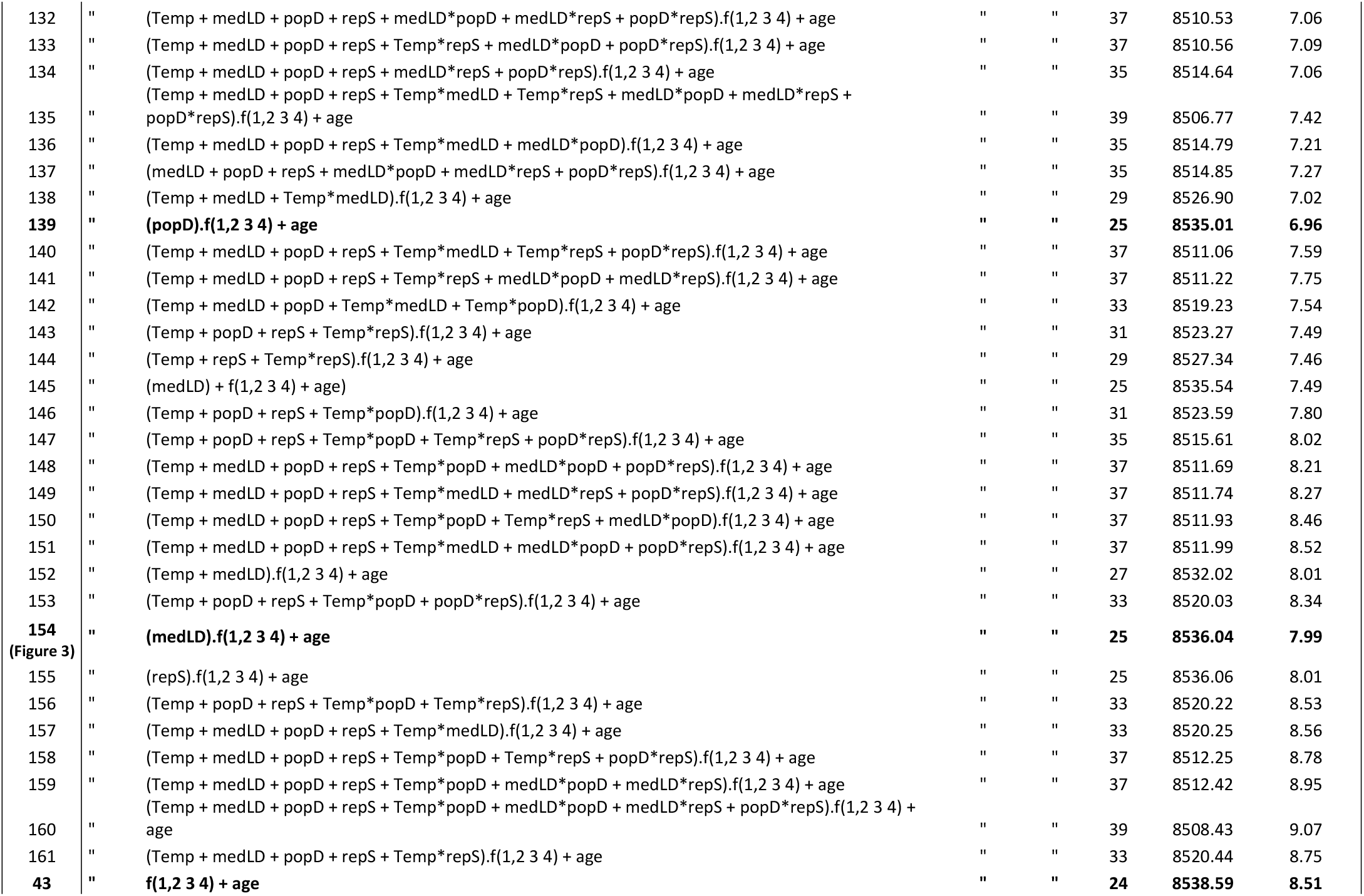
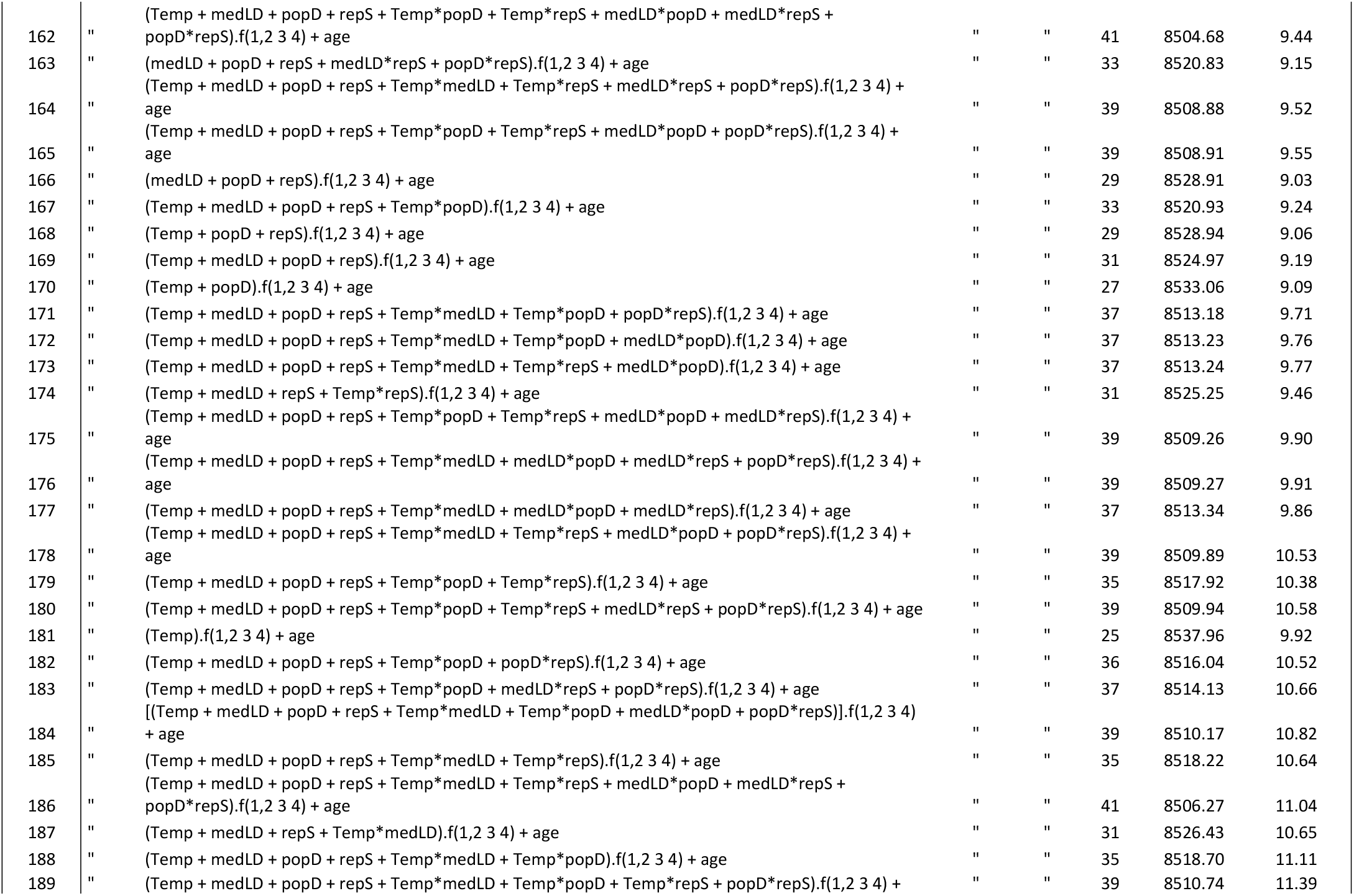
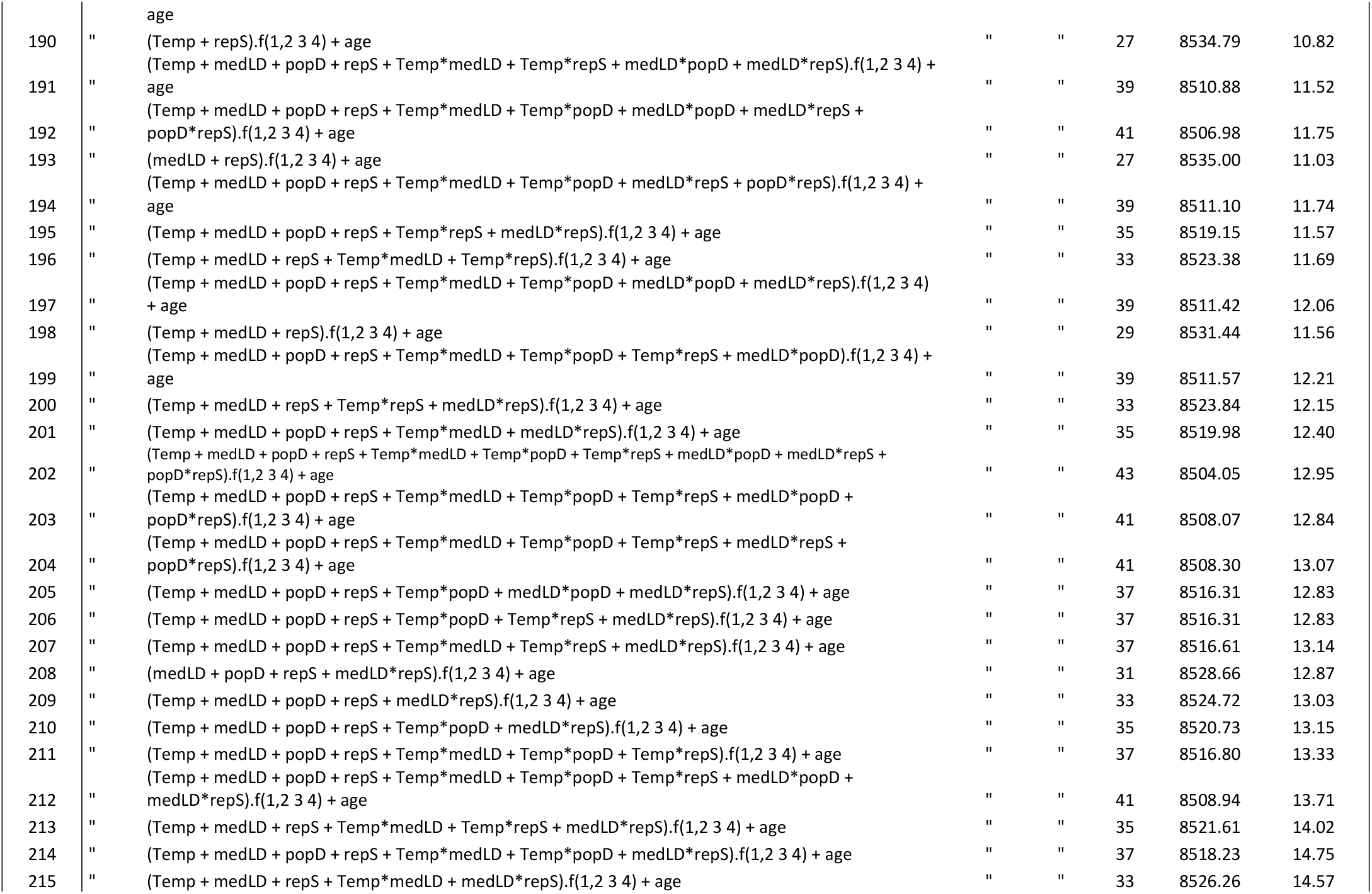
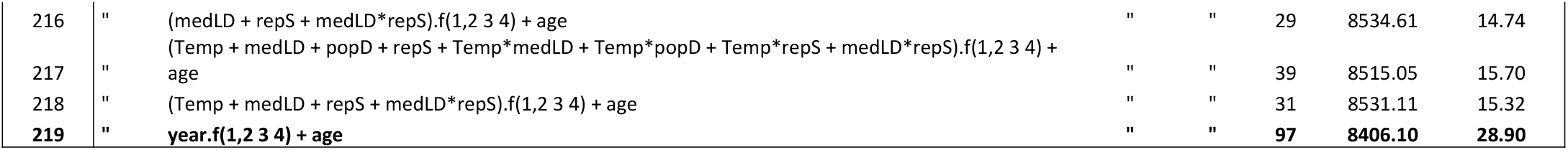
Multistate model selection assessing the effects of population-level annual covariates on the relative laying date - adult survival link. The effects of 4 annual covariates (Temp = mean daily mean temperature from March 31^st^ to May 7^th^, repS = reproductive success, popD = population density, medLD = median laying date) and their first-order interactions as continuous temporal covariates on survival probabilities are tested on one of the best models from the previous selection (Table 3): Model 43. Starting model: Model 43, best model: Model 103. AICc(Model 103) = 8578.5326. For initial state distribution: “to” means that at their first capture, individuals are not evenly distributed among the four phenological quartiles. For between states transition probabilities: “f.to” means that transition probabilities depend on the departure state (f), the arrival state (to) and their interaction. For survival: “f” means that survival depends on the current state of the individual. np: number of parameters.

**Table A5:**
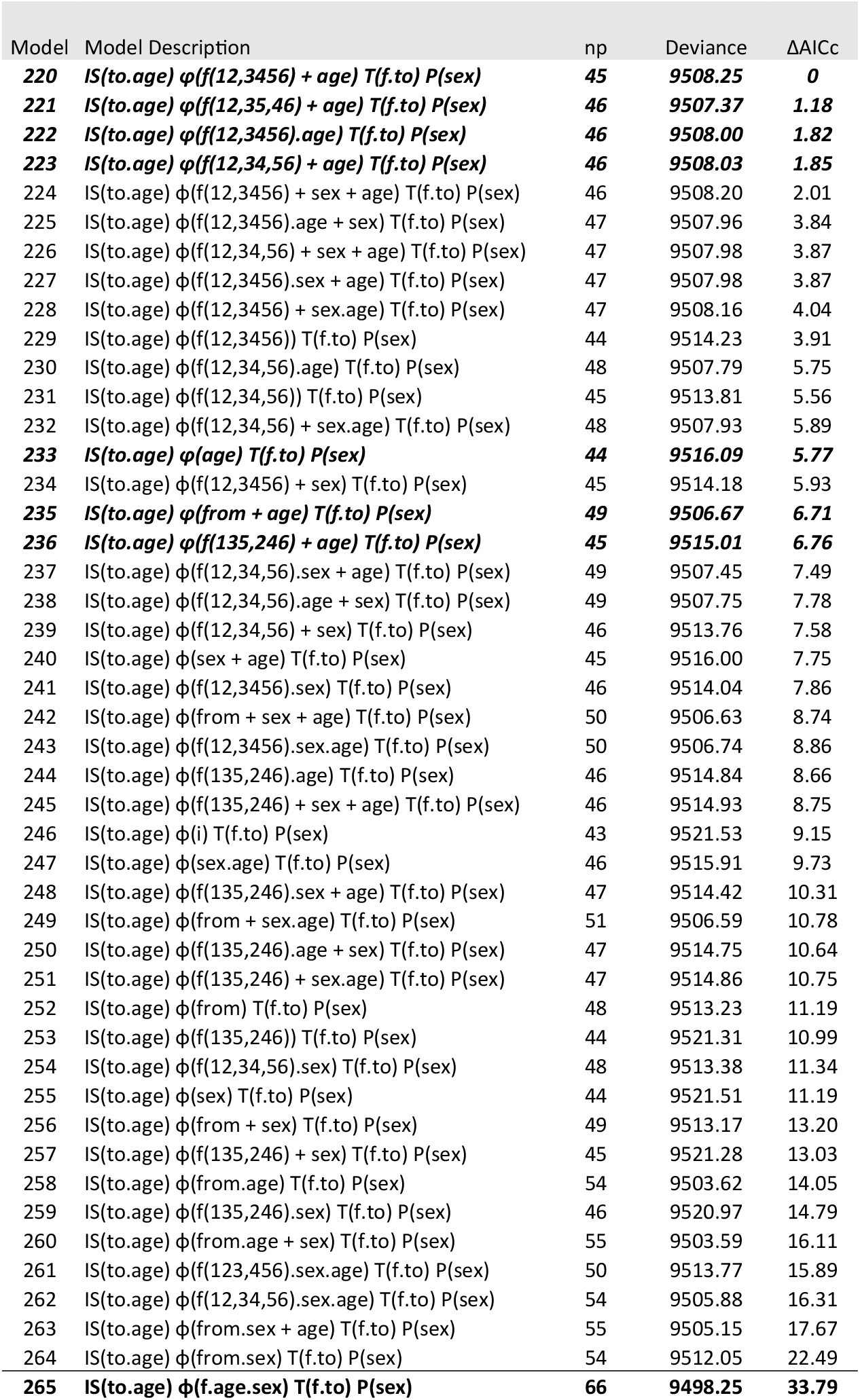
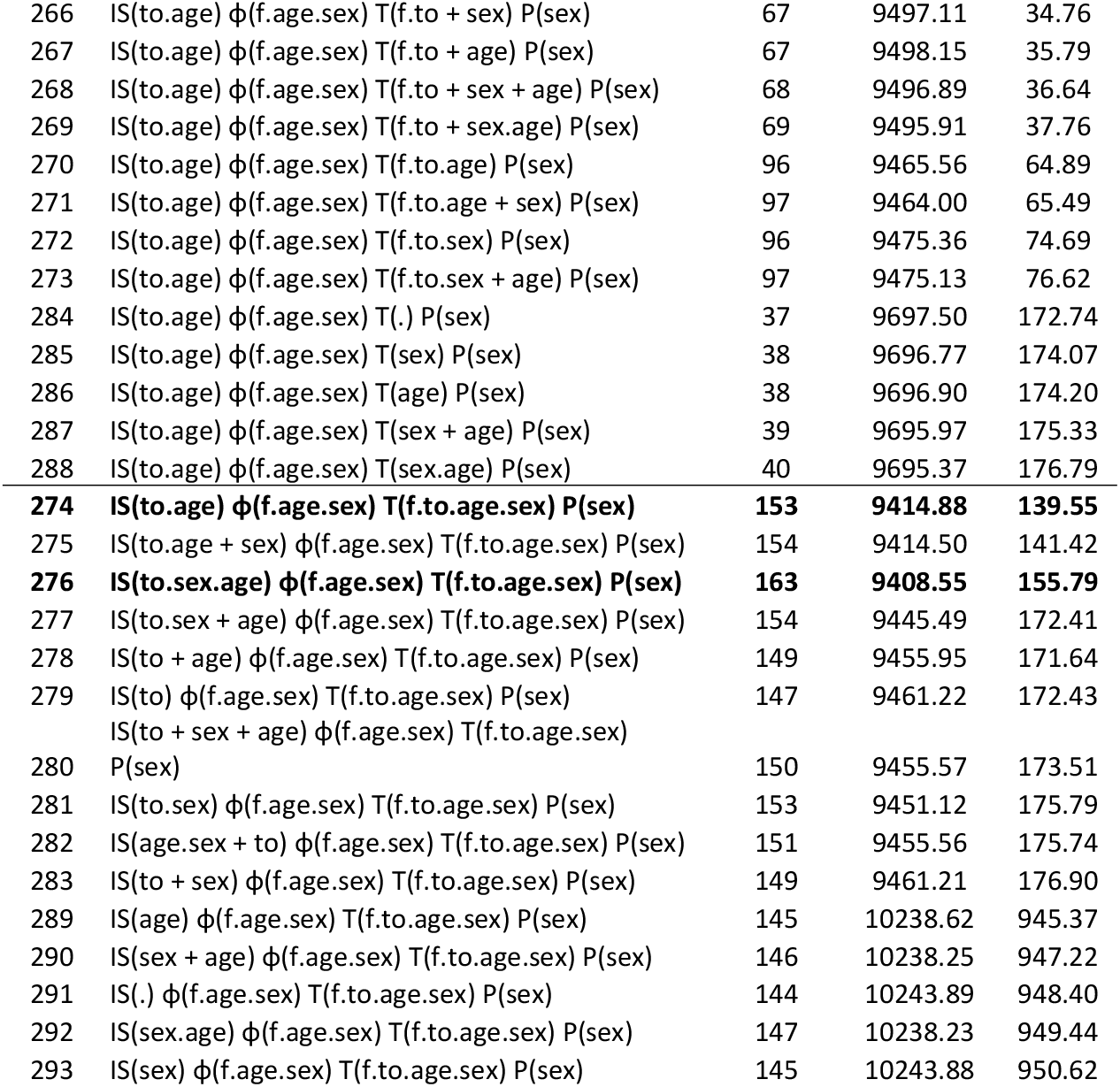
Multistate model selection assessing the links between individual breeding success, relative laying date and adult survival. The 6 states represent different combinations of individual breeding success and relative laying date (see Figure 5 for details). Starting model: Model 276 included the interactive effects of age, sex and state for its various structural components. Recapture probabilities were assumed to depend on sex only. Sequential selection procedure was the same as in the previous model selection (Table 4). Best model: Model 220. AICc(Model 220) = 9599.8097. np: number of parameters.

## Appendix 3: Temporal variation in survival in the two age classes

A series of additional models were made to check that the relationship between the survival of individuals and the variables presented in the results (median laying date medLD and population density popD) was not due to interannual variations in the proportion of age classes. (1 year old individuals 1yo versus 2 or more years 2yo). Additional survival models consider that the survival of a single age class varies over time while the other is constant (e.g., 2yo + 1yo.year) or that the survival of a single class is constrained by the median laying date or the population density (e.g., 2yo + 1yo.medLD). The results are presented in Table A6 below and illustrated in Figure A1.

**Table A6:** Monostate model selection assessing the link between the median laying date medLD, the population density and average annual survival. Best model: Model ϕ (age + popD) p(sex), AiCc = 2547.6513. p: number of parameters.

| Model description | np | Deviance | $\Delta AICc$ |
| --- | --- | --- | --- |
| $\phi$ (age + popD) p(sex) | 6 | 2535.62 | 0 |
| $\phi$ (age + medLD) p(sex) | 6 | 2536.86 | 1.24 |
| $\phi$ (1yo + 2yo.popD) p(sex) | 6 | 2537.76 | 2.14 |
| $\phi$ (2yo + 1yo.popD) p(sex) | 6 | 2537.99 | 2.38 |
| $\phi$ (1yo + 2yo.medLD) p(sex) | 6 | 2538.14 | 2.52 |
| $\phi$ (age) P(sex) | 5 | 2540.29 | 2.67 |
| $\phi$ (2yo + 1yo.medLD) p(sex) | 6 | 2538.91 | 3.30 |
| $\phi$ (1yo + 2yo.year) p(sex) | 41 | 2476.02 | 11.78 |
| $\phi$ (2yo + 1yo.year) p(sex) | 38 | 2504.29 | 33.85 |
| $\phi$ (age.year) p(sex) | 74 | 2440.68 | 45.63 |

**Figure A1:**
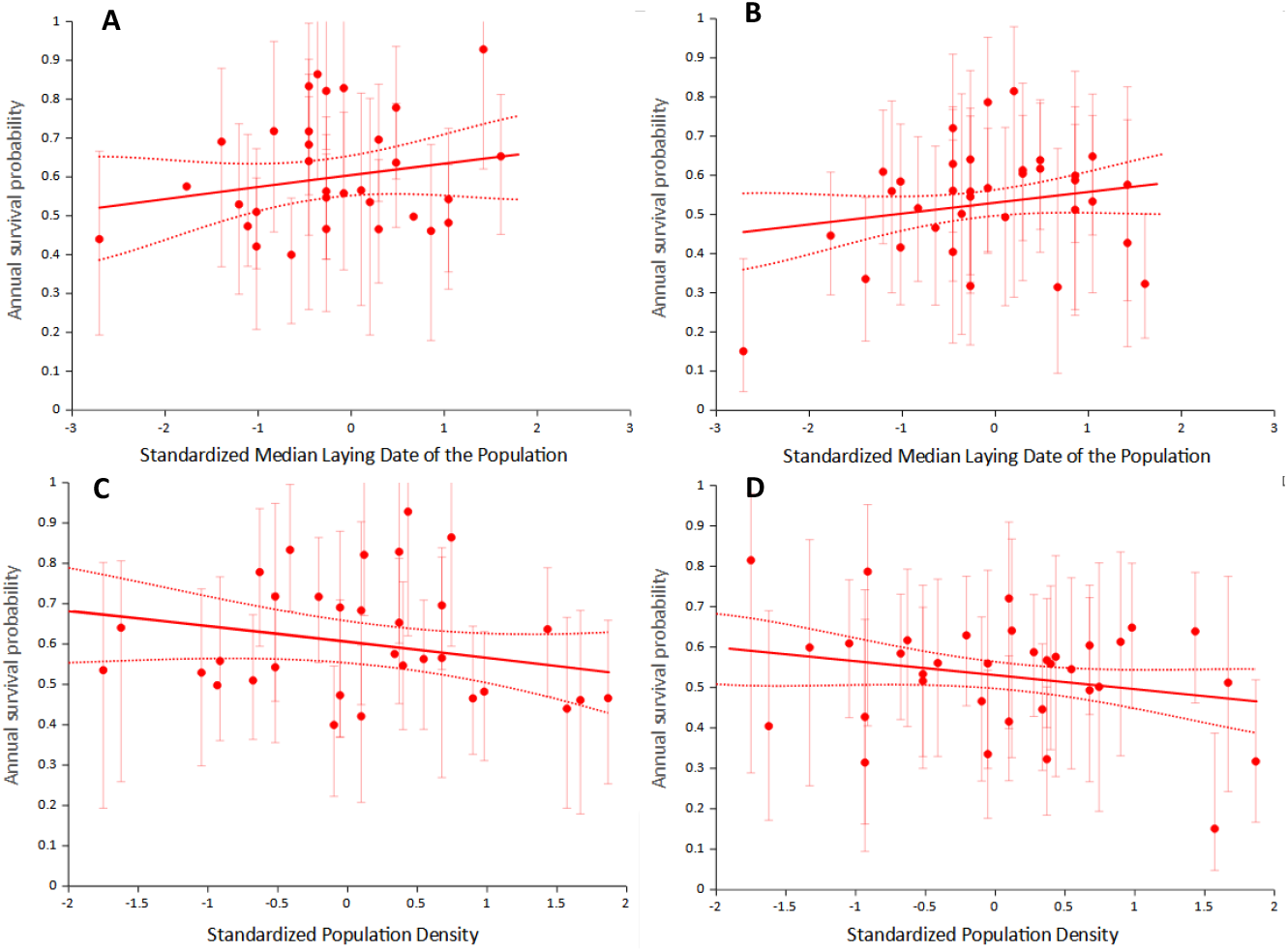
Relationship between annual survival probabilities and two demographic parameters. **A**: the survival probability of 1 year old individuals (1yo) is positively correlated with median laying date of the population in the previous spring (medLD). The points are the estimates of Model ϕ (2yo + 1yo.year) p(sex) in Table A6 for 1yo, with 95% confidence intervals, the solid line is for 1yo from Model ϕ (2yo + 1yo.medLD) p(sex) in Table A6, which has a continuous effect of medLD on survival, its 95% CI are represented by the dashed lines. **B**: the survival probability of 2 years old or older individuals (2yo) is positively correlated with medLD. The points are the estimates of Model ϕ (1yo + 2yo.year) p(sex) in Table A6 for 2yo, with 95% confidence intervals, the solid line is for 2yo from Model ϕ (1yo + 2yo.medLD) p(sex) in Table A6, which has a continuous effect of medLD on survival, its 95% CI are represented by the dashed lines. **C**: the survival probability of 1yo is negatively correlated with population density during the previous spring (popD). The points are the estimates of Model ϕ (2yo + 1yo.year) p(sex) in Table A6 for 1yo, with 95% confidence intervals, the solid line is for 1yo from Model ϕ (2yo + 1yo.popD) p(sex) in Table A6, which has a continuous effect of popD on survival, its 95% CI are represented by the dashed lines. **D**: the survival probability of 2yo is positively correlated with popD. The points are the estimates of Model ϕ (1yo + 2yo.year) p(sex) in Table A6 for 2yo, with 95% confidence intervals, the solid line is for 2yo from Model ϕ (1yo + 2yo.popD) p(sex) in Table A6, which has a continuous effect of popD on survival, its 95% CI are represented by the dashed lines.

## Appendix 4: Repeatability of laying date percentile in the E-Pirio population

We performed a repeatability analysis based on 1000 parametric bootstraps as implemented in the rptR package (Stoffel et al., 2018) of the R-software v3.5.1 (R Core Team, 2018). The statistical significance of the repeatability of each metric was tested by a likelihood ratio test comparing the model fit of a model including grouping factor (here, the species or the street) and one excluding it.

A significant repeatability was found for both males and females:

For males: 712 observations, 382 individuals, R = 0.124 [0.027; 0.220]_95%CI_, likelihood ratio test: D=8.51, df=1, P=0.02

For females: 818 observations, 410 individuals, R = 0.406 [0.316; 0.493]_95%CI_, likelihood ratio test: D=95.1, df=1, P=0

## Appendix 5: Matrix of probabilities of transition among the different relative laying date states as described in Table 3 and Figure 2

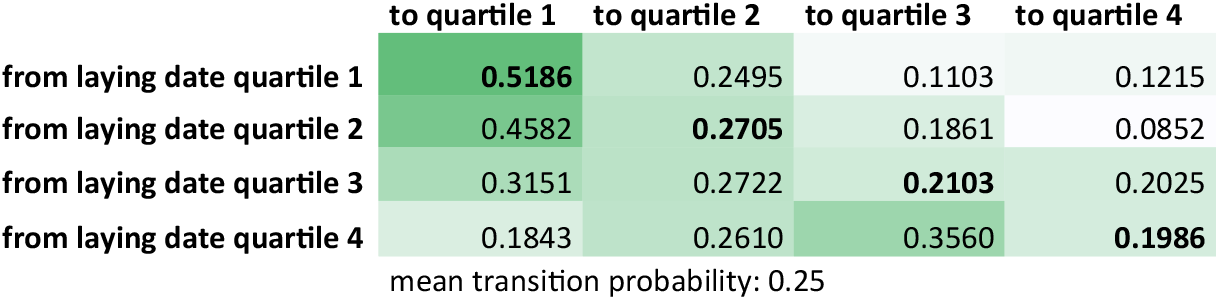

Quartile 1 corresponds to the 25% earliest breeders of the population in a given year, quartile 4 to the latest. From all starting laying date quartiles, the probability of transitioning to the same or an earlier quartile are higher than transitioning to a later quartile.

