## Supplementary material for "Is earlier reproduction associated with higher or lower survival? Antagonistic results between individual and population scales in the blue tit": Data description file

CMR DATA

1) Monostate_CMR_histories
first column: CMR histories 1979-2016
second column: right censoring
third column: sex (1: male; 2: female)
fourth column: age class at first capture (1: 1 year old; 2: 2+ years old)


2) Multistate_LD_CMR_histories
first column: CMR histories 1979-2016
Numbers ranging from 1 to 4 correspond to the quartile of the focal individual within the laying date distribution of the population in a given year
second column: right censoring
third column: sex (1: male; 2: female)
fourth column: age class at first capture (1: 1 year old; 2: 2+ years old)


3) Multistate_LD_repS_CMR_histories
first column: CMR histories 1979-2016
Numbers ranging from 1 to 6 correspond to the combination of relative laying date and fledgling success of the focal individual as described in Fig5A of the main document
second column: right censoring
third column: sex (1: male; 2: female)
fourth column: age class at first capture (1: 1 year old; 2: 2+ years old)


TEMPORAL COVARIATES (Temporal_covariates.txt)

popD: normalized population density in year t
medLD: median laying date in the population in year t+1
repS: mean number of fledglings per attempted first brood in the population in year t+1
