## Supplementary material for "Is earlier reproduction associated with higher or lower survival? Antagonistic results between individual and population scales in the blue tit": model description file

General
in all models' names:
- phi : survival probability
- p : recapture probability
- t : transition probability among laying date or breeding success states
- i : initial state
- t : time (year effect)
- a : age class
- captage : group of age class at first capture
- s : sex
- full : structure of the starting models for unspecified parameters
- CI : computation of confidence intervals


Monostate_models_Simple
This set of models includes all monostate models without temporal covariates


Monostate_models_Covariates
This set of models includes all monostate models with temporal covariates
popD: normalized population density in year t
medLD: median laying date in the population in year t+1
repS: mean number of fledglings per attempted first brood in the population in year t+1


Multistate_LD
This set of models includes all multistate models with laying date states, without temporal covariates
Numbers ranging from 1 to 4 correspond to the quartile of the focal individual within the laying date distribution of the population in a given year


Multistate_LD_Covariates
This set of models includes all multistate models with laying date states, with temporal covariates
Numbers ranging from 1 to 4 (brackets following "f") correspond to the quartile of the focal individual within the laying date distribution of the population in a given year
Numbers ranging from 1 to 10 (brackets following "t") correspond to the temporal covariates included and their interactions, based on the following code:
1 : temperature
2 : laying date
3 : population density
4 : fldgling success
5 : 1*2
6 : 1*3
7 : 1*4
8 : 2*3
9 : 2*4
10 : 3*4


Multistate_LD_repS
This set of models includes all multistate models with both laying date and success states (no temporal covariates)
Numbers ranging from 1 to 6 correspond to the combination of relative laying date and fledgling success of the focal individual as described in Fig5A of the main document
